# Molecular insights into atypical modes of β-arrestin interaction with seven transmembrane receptors

**DOI:** 10.1101/2023.07.05.547776

**Authors:** Jagannath Maharana, Fumiya K. Sano, Parishmita Sarma, Manish K. Yadav, Longhan Duan, Tomasz M. Stepniewski, Madhu Chaturvedi, Ashutosh Ranjan, Vinay Singh, Sayantan Saha, Gargi Mahajan, Mohamed Chami, Wataru Shihoya, Jana Selent, Ka Young Chung, Ramanuj Banerjee, Osamu Nureki, Arun K. Shukla

## Abstract

β-arrestins are multifunctional proteins that are critically involved in regulating spatio-temporal aspects of GPCR signaling. The interaction of β-arrestins with GPCRs is typically conceptualized in terms of receptor activation and phosphorylation primarily in the carboxyl-terminus. Interestingly however, there are several GPCRs that harbor majority of phosphorylation sites in their 3^rd^ intracellular loop (ICL3) instead of carboxyl-terminus but still robustly engage β-arrestins. Moreover, there are several 7TMRs that are now characterized as intrinsically-biased, β-arrestin-coupled receptors (ACRs) due to lack of functional G-protein-coupling but robust β-arrestin binding leading to functional outcomes. The molecular basis of β-arrestin interaction and activation upon binding to these types of 7TMRs is currently elusive, and it represents a major knowledge gap in our current understanding of this signaling system. Here, we present seven cryo-EM structures of β-arrestins in basal state, activated by the muscarinic M2 receptor (M2R) through its ICL3, and a β-arrestin-coupled receptor known as decoy D6 receptor (D6R). These structural snapshots combined with biochemical, cellular, and biophysical experiments including HDX-MS and MD simulation provide novel insights into the ability of β-arrestins to preferentially select specific phosphorylation patterns in the receptors, and also illuminate the structural diversity in 7TMR-β-arrestin interaction. Surprisingly, we also observe that the carboxyl-terminus of β-arrestin2 but not β-arrestin1 undergoes structural transition from a β-strand to α-helix upon activation by D6R, which may preclude the core-interaction with the activated receptor. Taken together, our study elucidates previously unappreciated aspects of 7TMR-β-arrestin interaction, and provides important mechanistic clues about how the two isoforms of β-arrestins can recognize and regulate a large repertoire of GPCRs.

## Introduction

β-arrestins (βarrs) are multifunctional proteins that interact with, and regulate a large repertoire of G protein-coupled receptors (GPCRs) at multiple levels (*1–4*). The interaction of GPCRs and βarrs is typically conceived to be driven primarily by agonist-induced receptor phosphorylation and receptor activation although emerging studies have started to suggest additional contributing factors such as membrane interaction, catalytic activation, and role of specific phospholipids (*2–10*). A number of structures of GPCR-βarr1 complexes have been determined in the past couple of years, which have provided the first glimpse of high-resolution information about this interaction (*11–16*). Still however, considering the divergent primary sequence and phosphorylation patterns of GPCRs, the molecular mechanisms driving the broadly conserved nature of GPCR-βarr interaction and activation remains elusive to a large extent until recently. Some recent studies however have started to shed light on phosphorylation-mediated components of GPCR-βarr binding through broadly conserved phosphorylation motifs identified in a large number of GPCRs (*17–20*). For example, structural and biophysical studies have proposed the framework of phosphorylation codes and modulatory sites in the GPCR carboxyl-terminus as a possible mechanism governing phosphorylation-mediated βarr interaction (*19, 20*). More recently, two independent structural studies have identified that the presence of a P-X-P-P type phosphorylation motif in the carboxyl-terminus of a broad set of GPCRs, where P is a phosphorylation site, is a critical determinant of βarr interaction and activation (*17, 18*). Still however, there are several key questions about this versatile interaction that remain unanswered and represent important knowledge gaps in our current understanding of this signaling and regulatory paradigm.

There are several GPCRs, for example the human muscarinic receptor subtype 2 (M2R), that contain a short carboxyl-terminus with a very few potential phosphorylation sites, but they harbor phosphorylation sites primarily in their 3^rd^ intracellular loop (*5, 21–24*). Site-directed mutagenesis and biochemical studies have demonstrated the contribution of phosphorylation sites in the intracellular loops of some of these receptors to contribute towards βarr binding (*25, 26*). However, whether these receptors engage the same binding interface with βarrs and impart similar activation features as other GPCRs with phosphorylation sites on their carboxyl-terminus remains primarily unexplored in terms of direct structural visualization. Moreover, there are several 7TMRs such as the human decoy D6 receptor (D6R), sometimes classified as non-signaling or non-functional GPCRs as they lack functional G-protein-coupling, but robustly interact with, and signal through βarrs (*27–30*). The molecular mechanisms engaged by these receptors, known as atypical chemokine receptors (ACKRs) or Arrestin-coupled receptors (ACRs), to bind and activate βarrs are also mostly elusive with respect to the binding interface and activation dependent conformational changes vis-à-vis prototypical GPCRs (*31–34*). The paucity of structural information and functional correlation on β-arrestin interaction and activation by the ACRs, and GPCRs engaging βarrs through their ICL3, limits current understanding of structural and functional diversity encoded in the 7TMR-β-arrestin system.

Accordingly, here we visualize the structural details of βarr interaction and activation by M2R and the D6R using cryogenic-electron microscopy (cryo-EM). The structural snapshots of M2R-βarr complexes uncover the precise interaction interface between ICL3 and βarrs for the first time. Surprisingly, we observe an α-helical conformation adopted by the distal carboxyl-terminus of βarr2 but not βarr1, upon activation by the phosphorylated D6R. We complement the key findings uncovered by the structural snapshots with HDX-MS, molecular dynamics simulation, and cellular assays. Taken together, our findings provide previously lacking and unanticipated aspects of 7TMR-βarr interaction and activation, and significantly advance the current conceptual framework in the field with direct implications for exploring novel therapeutic avenues.

## Results

In order to visualize the atypical modes of βarr recruitment, we focused our efforts on the M2R which has a short carboxyl-terminus with the majority of potential phosphorylation sites localized in ICL3, and D6R that is intrinsically βarr-biased receptor with no detectable G-protein activation despite robust βarr binding and signaling (Figure 1). We used full-length, wild-type M2R phosphorylated *in-cellulo* via co-expression of a membrane-tethered GRK2 construct (GRK2^CAAX^) and agonist-induced phosphorylation followed by incubation with purified βarr1 and Fab30 to reconstitute the complex (Figure S1A-B). Subsequently, we attempted to determine the structure of this complex using cryo-EM, and while the receptor component was not resolved at high-resolution, presumably due to inherent flexibility, we successfully determined the structure of receptor-bound βarr1 at 3.1Å resolution with focused refinement (Figure 1C and Figure S2). In order to reduce the flexibility of the receptor component in this complex, we cross-linked the pre-formed M2R-βarr1-Fab30 complex using on-column glutaraldehyde cross-linking (*35*) followed by cryo-EM data collection. Still however, the receptor exhibited flexible positioning relative to βarr1, and therefore, we could determine the structure of only the receptor-bound βarr1 at 3.2Å (Figure 1C, Figure S1C-D, and Figure S3). Nonetheless, these structural snapshots allowed us to identify the phosphorylated region of the ICL3 in M2R that forms the key interaction interface with βarr1, and thereby allowed us to synthesize and validate the corresponding phosphopeptide (M2Rpp) (Figure S4A-B), and determine the structure of M2Rpp-βarr2-Fab30 complex at 2.9Å resolution (Figure 1C, Figure S1E-F, and Figure S5).

**Fig. 1.**
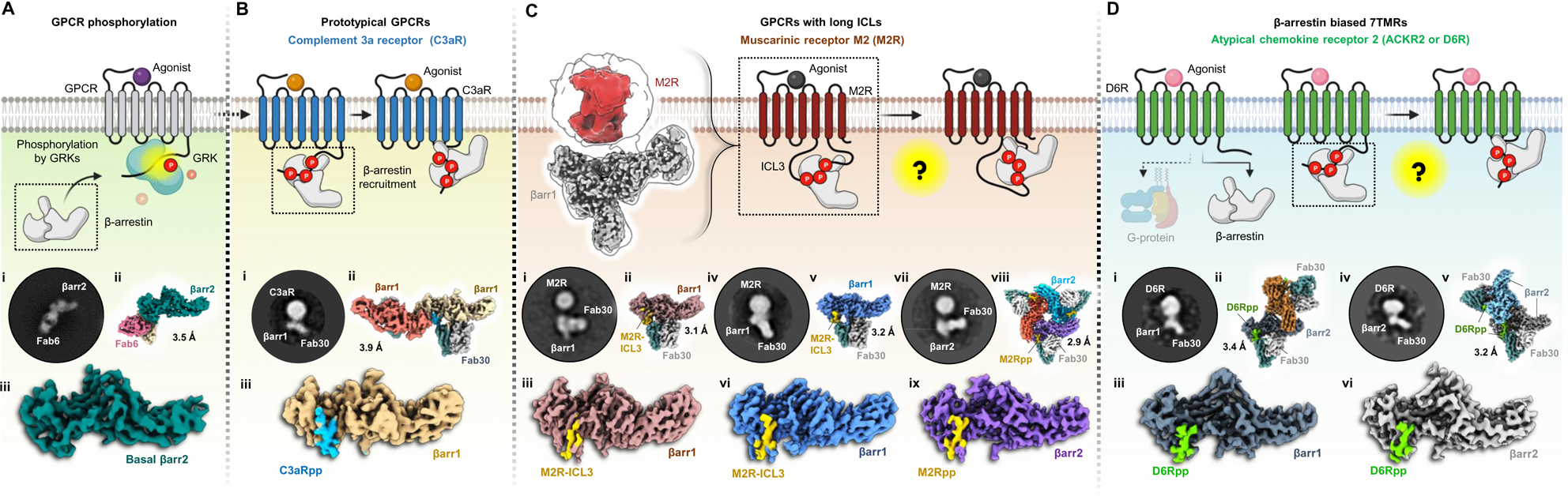
A structural approach to understand the atypical modes of βarr interaction with 7TMRs. **(A)** Phosphorylation by GRKs mediate βarr interaction with GPCRs. Cryo-EM structure of full-length βarr2 sheds light into its basal state conformation. (**i, ii, iii**) 2D class average, overall 3D map of βarr2 bound to Fab6 and structure of βarr2 alone are shown. **(B)** Two distinct modes of interactions of β-arrestins with the phosphorylated tail of GPCRs. The phosphorylation pattern of complement receptor C3aR has been utilized to delineate the “hanging” mode of βarr interaction. (**i, ii, iii**) 2D class average, overall dimeric 3D map and structure of C3aRpp-βarr1 are presented. **(C)** Muscarinic receptor 2 was chosen to represent the class of GPCRs that control βarr activation and signalling through extended intracellular loops. Lack of phosphorylation at the C-terminus raises the question of the existence of the biphasic mode of βarr interaction. A 3D reconstruction has been shown to the left to show a “hanging” mode of complex organization. High resolution structures of M2R-ICL3 bound βarr1/2 are shown below. 2D class average, overall 3D map and structure of (**i, ii, iii**) M2R-βarr1, (**iv, v, vi**) M2R-βarr1 of cross-linked complex, and (**vii, viii, ix**) M2Rpp-βarr2 are shown. **(D)** βarr biased 7TMRs lack G protein coupling, but signal through βarrs. The mode of βarr interaction to this class of receptors is yet to be explored and presented as a schematic diagram. To explore the possibilities of the “hanging” mode, structures of βarr1/2 have been determined in complex with the phosphorylation pattens of the Decoy receptor D6R or ACKR2. 2D class average, overall dimeric 3D map and structure of (**i, ii, iii**) D6Rpp-βarr1, and (**iv, v, vi**) D6Rpp-βarr2 have been shown. The estimated resolutions for all the structures have been mentioned against each map.

For D6R, we have reported previously that the critical determinants of βarr recruitment are located primarily in its carboxyl-terminus (*28*), and therefore, we generated a set of phosphopeptides corresponding to the phosphorylated D6R and tested their ability to activate βarrs *in-vitro* using Fab30 reactivity or limited proteolysis as readouts (Figure S4C-F). Based on these assays, we identified D6Rpp2, referred to as D6Rpp from here onwards, to activate βarrs most efficiently, and we used it to reconstitute D6Rpp-βarr1/2-Fab30 complexes (Figure S1G-H), and determined their structures at 3.4Å and 3.2Å resolution, respectively (Figure 1D, Figure S6, and Figure S7). In addition, we also determined the structures of wild-type βarr2 in its basal conformation stabilized by Fab6 (Figure 1A, Figure S1I, and Figure S8), and βarr1 in complex with a carboxyl-terminus phosphopeptide of the complement C3a receptor (C3aR), a prototypical GPCR (Figure 1B, Figure S1J, and Figure S9), as references for basal and typical active conformations. The electron densities of the phosphorylated receptor domains and the key loops in βarrs in these above-mentioned structures are presented in Figure S10.

The M2R-βarr1-Fab30 complexes reminisce a hanging conformation observed previously for prototypical GPCRs (*11, 35*) with a significant spacing between the receptor and βarr components, presumably due to their interaction mediated primarily through the long ICL3 (∼150 residues) in the M2R (Figure 2A-E). Not only this is observed in M2R complexes with both isoforms of βarrs but also in complexes where the receptor is phosphorylated by either GRK2 or GRK6 (Figure 2A), suggesting that hanging conformations represent a significant population in M2R-βarr interaction irrespective of βarr or GRK isoforms. While glutaraldehyde cross-linking appears to stabilize a more closely engaged complex as reflected in negative-staining 2D class averages (Figure 2G), it did not significantly help resolve the receptor component better compared to the non-cross-linked complex in cryo-EM. The structure of M2R-bound βarr1 revealed a phosphorylated stretch of ICL3 in the receptor that harbors the residues from Q^303^-G^313^ with four phosphorylation sites (Thr^308^, Ser^310^, Thr^311^ and Ser^312^), and it docks on the N-domain of βarr1 (Figure 2F and Figure 2H). Interestingly, M2Rpp that is derived from the ICL3 sequence visualized in M2R-bound βarr1 structure binds to an analogous interface on βarr2 (Figure 2I-J). The βarr1 and 2 in these structures exhibit an inter-domain rotation of ∼18° and 23°, respectively, disruption of the three-element and polar-core network (Figure S11A-D), and significant reorientation of the critical loops compared to the basal conformation (Figure S11E). Notably, the phosphate groups in the M2R-ICL3 stretch resolved in these structures are organized in a P-X-P-P pattern, where P is a phosphorylation site, and they are engaged in ionic interactions with conserved Lys and Arg residues in βarrs organized in K-R-K type pattern involving Arg^7/8^, K^10/11^, K^11/12^, R^25/26^, K^107/108^ and K^294/295^ (Figure 2K). A comprehensive list of residue-residue contacts between the phosphopeptides and βarrs have been provided in Table S3.

**Fig. 2.**
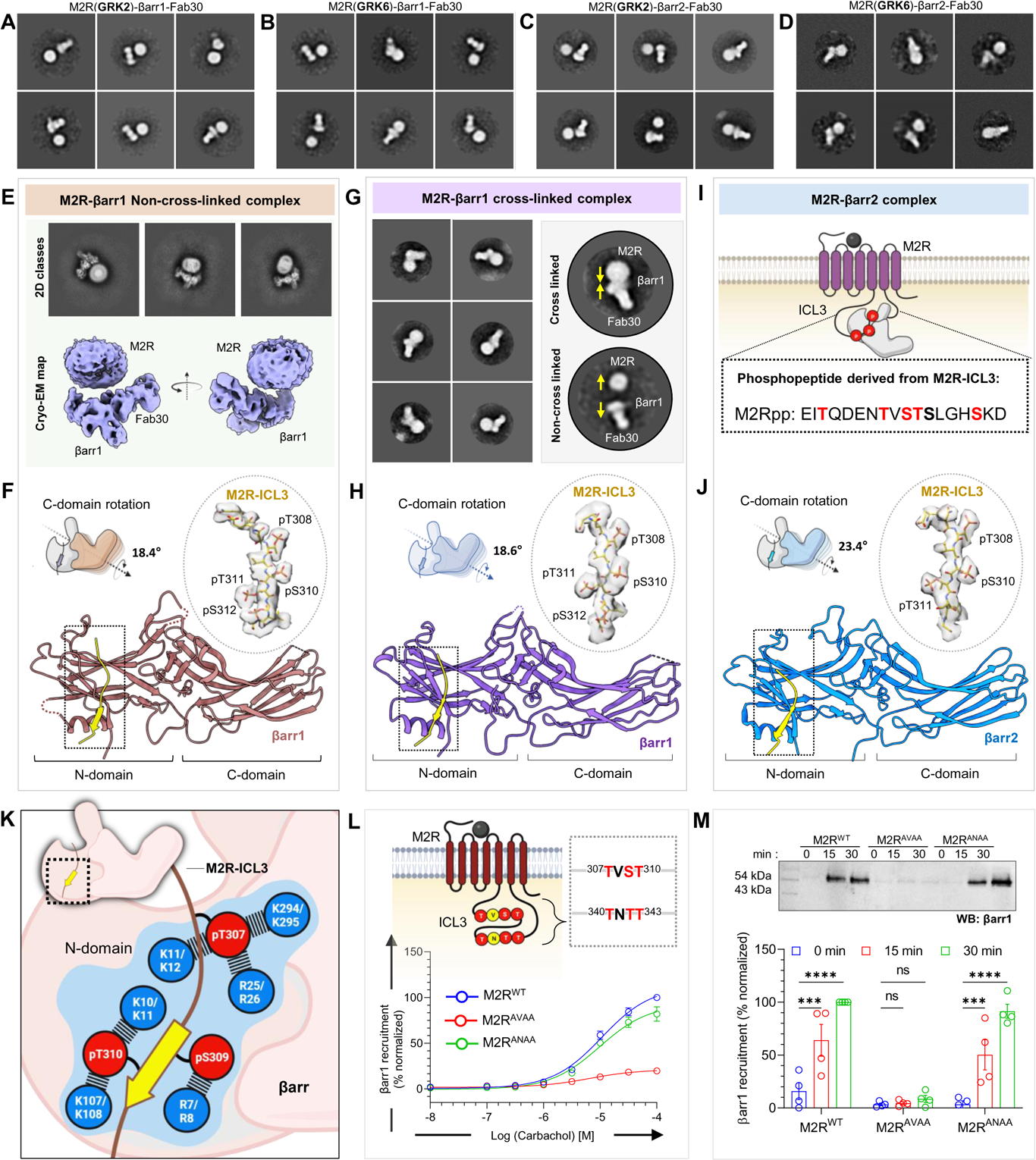
Structural insights into ICL3 driven βarr interaction with M2R. **(A, B, C, D)** Negative staining EM class averages of M2R, endogenously phosphorylated by GRK2/6 in complex with βarr1 or βarr2. **(E)** Cryo-EM 2D classes, 3D reconstruction of “hanging” M2R-βarr1-Fab30 complex. **(F)** structure of βarr1 bound to phosphorylated M2R-ICL3. The EM density of ICL3 has been shown in inset. βarr1 attains an active conformation with a C-domain rotation of 18.4° with respect to the N-domain. **(G)** On-column crosslinking was performed to rigidify the M2R-βarr1 complex. Representative negative staining EM 2D classes have been presented to depict the effect of cross-linking. While yellow arrows show potential movement of the complex subunits. **(H)** Structure of cross-linked M2R-βarr1 complex. The EM density of ICL3 has been shown in inset. C-domain rotation value with respect to N-domain is 18.6°. **(I)** Sequence of phosphopeptide derived from the ICL3 of M2R. **(J)** Structure of M2Rpp-βarr2 in ribbon representation. M2Rpp is shown in yellow and βarr2 in blue. Density map of phosphopeptide has been displayed to the left. βarr2 attains an active conformation with 23.4° rotation of C-domain upon activation with M2Rpp. **(K)** The phosphorylated residues from ICL3 make critical contacts with the Lys and Arg residues present on the N-domain of βarrs. Lys and Arg residues of βarr1 (upper) and βarr2 (lower) have been highlighted in blue. **(L)** Cartoon representation illustrating the presence of possible phosphorylation clusters in the ICL3 of M2R. Mutations of the two phosphor-motifs: TVST and TNTT have been generated to assess the βarr recruitment measured by bystander NanoBiT assay (receptor+SmBiT-βarr1+LgBiT-CAAX). Substitution of phosphosites of TVST to AVAA leads to abrupt reduction in βarr recruitment, whereas, TNTT to ANAA substitution maintained βarr recruitment, suggesting critical role played by TVST on βarr recruitment to M2R. (mean±SEM; n=3; normalized with respect to highest ligand concentration signal for M2RWT as 100%). (**M**) Role of TVST in βarr recruitment is further corroborated by co-immunoprecipitation assay. On Carbachol stimulation, M2RAVAA showed dramatic reduction in βarr1 recruitment. A representative blot and densitometry-based quantification (mean±SEM; n=4; normalized with M2R 30min stimulation condition signal as 100%; Two-way ANOVA, Tukey’s multiple comparisons test) is presented. The exact p values are as follows: M2R^WT^ - 0 vs. 15min = 0.0006, M2R^WT^ - 0min vs. 30min = <0.0001, M2RANAA - 0min vs. 15min = 0.0008, M2RANAA - 0min vs. 30min = <0.0001.(***p = 0.0001; ****p < 0.0001; ns, non-significant).

The sequence analysis of M2R reveals that there are two plausible P-X-P-P type motifs in the ICL3, one represented by T^308^-V-S^310^-T^311^ that is observed in the structures presented here while the other is represented by T^340^-N-T^342^-T^343^ (Figure 2L). Therefore, in order to further validate the key contribution of T-V-S-T stretch in M2R-ICL3 in βarr engagement and activation, we generated two different mutants of the receptor with the phosphorylation sites in each of these P-X-P-P motifs changed to Ala residues by site-directed mutagenesis. Subsequently, we measured agonist-induced βarr1 recruitment to these mutants vis-à-vis the wild-type receptor using NanoBiT and co-immunoprecipitation assay, and observed that mutation of T-V-S-T, but not T-N-T-T, nearly ablates βarr binding (Figure 2L-M and Figure S12). These observations establish the key contribution of the T-V-S-T motif in M2R-ICL3 in driving βarr recruitment, and also underscore the shared mechanism of βarr activation by M2R and other prototypical GPCRs despite distinct receptor domains engaging βarrs.

In contrast to prototypical GPCRs, some chemokine receptors such as CXCR7 and D6R, and a complement C5 receptor (C5aR2), lack G-protein-coupling but maintain robust βarr recruitment and downstream signaling (*28, 36–39*). These receptors, referred to as atypical chemokine receptors (ACKRs) or Arrestin-coupled Receptors (ACRs), are essentially intrinsically βarr-biased and represent an excellent model system to probe structural and functional diversity of βarrs. Thus, we next attempted to reconstitute D6R-βarr complexes using co-expression of the receptor, GRK2 or GRK6, and βarr1/2, followed by *in-cellulo* assembly of the complex via agonist-stimulation and stabilization using Fab30. While we observed clear complex formation and a typical architecture by negative staining that is reminiscent of the hanging conformation (Figure 3A-B), attempts to scale-up the complex for cryo-EM analysis were not successful. Therefore, we focused our efforts to determine the structures of βarrs in complex with a phosphorylated peptide corresponding to the carboxyl-terminus of D6R (D6Rpp). We first confirmed that D6R-βarr interaction depends on receptor phosphorylation by truncating the carboxyl-terminus of D6R harboring the phosphorylation sites, which resulted in near-complete ablation of agonist-induced βarr1 recruitment (Figure 3C). Subsequently, we characterized D6Rpp using *in-vitro* proteolysis and Fab30 reactivity assays (Figures S4C-F), and further validated βarr activation by this peptide using HDX-MS (Figure 3D). We observed that D6Rpp binding resulted in robust activation of βarrs as reflected by significant conformational changes in multiple β-strands and loop regions in the N-domain (Figure 3E-F and Figure S13). Interestingly, we also observed notable differences between the HDX-MS pattern of βarr1 vs. βarr2 such as reduced solvent exposure of β-strand XIV and XV in the C-domain of βarr2, which suggests isoform-specific differences between activation of βarr1 vs. βarr2.

**Fig. 3.**
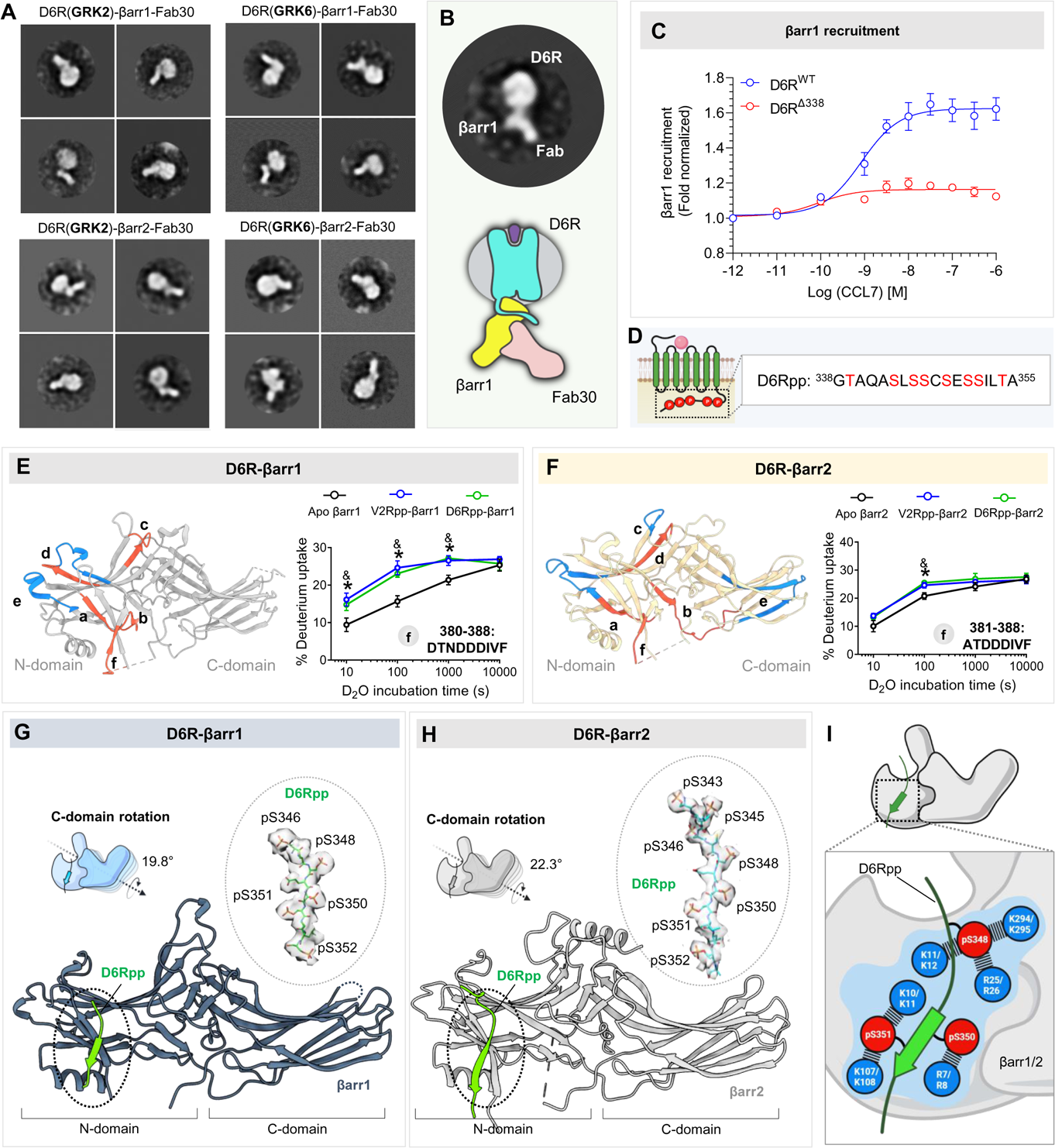
Structural insights into D6R-βarr complex interaction and activation. **(A)** Reconstitution of D6R-βarr1 complex and visualization by negative staining EM. 2D class averages of D6R-βarr1/2 complexes endogenously phosphorylated with GRK2/6**. (B)** A representative 2D class average has been illustrated to highlight the “hanging” mode of βarr1 interaction with the receptor. **(C)** Dose response curve for CCL7-induced βarr1 recruitment for the mentioned D6R constructs using NanoBiT assay (Receptor-SmBiT+LgBiT-βarr1) (mean±SEM; n=3; normalized with respect to the lowest ligand concentration signal as 1). **(D)** Design of selected phosphopeptide derived from the C-terminus of D6R. (**E, F**) HDX-MS plots to show the potential of generated phosphopeptides from D6R to activate βarr1 and βarr2, respectively. Among regions (a-f) showing significant changes upon deuterium exchange, the fragment at the C-terminus (f) has been demonstrated to show activation of βarrs upon D6Rpp binding. (**G**) Structure of D6Rpp-βarr1 complex in ribbon representation. The density map of D6Rpp has been shown to the left. C-domain rotation of βarr1 bound to D6Rpp is 19.8°. (**H**) Structure of D6Rpp-βarr2 complex in ribbon representation. The density map of D6Rpp has been shown to the left. C-domain rotation of βarr2 bound to D6Rpp was calculated to be 22.3°. (**I**) The phosphorylation pattern from D6Rpp engage with a network of Lys and Arg residues present on the N-domains of βarrs. Residues highlighted in blue circles show the Lys and Arg residues in βarr1 (upper) and βarr2 (lower) respectively.

Next, we determined the structures of βarr1 and βarr2 in complex with D6Rpp, stabilized by Fab30, at resolution of 3.4Å and 3.2Å, respectively (Figure 1D, Figure 3G-H). We observed a similar interaction interface of D6Rpp on N-domains of βarr1 and 2 although seven phosphates were resolved in βarr2 structure compared to five in βarr1 (Figure 3G-H). Interestingly however, we observed that three phosphate groups namely Ser^348^, Ser^350^ and Ser^351^ organized in a P-X-P-P pattern are engaged in most extensive interactions with selected Lys and Arg residues in the N-domain of βarrs (Figure 3I). Similar to M2R, there are two putative P-X-P-P motifs in D6Rpp as well, still however, our structural snapshots reveal that βarrs prefers one of them (Figure S14). Expectedly, we also observed significant inter-domain movement in D6Rpp-bound βarrs, the reorientation of the key loop regions compared to the basal state, and disruption of the three-element and polar core network (Figure S11). A comprehensive list of residue-residue contacts between the phosphopeptides and βarrs have been given in Table S3.

Surprisingly, the distal carboxyl-terminus of βarr2 (Tyr^391^-Lys^408^) in D6Rpp-bound conformation adopts an α-helical structure, which is positioned in the central crest of βarr2 (Fig. 4A-B) and makes extensive interactions (Figure S15). This α-helix in βarr2 forms a key dimerization interface for the two protomers in this structure and arranged in an anti-parallel coiled coil fashion with extensive contacts across the two protomers (Figure 4C, Table S4). We further analyzed the stability of this α-helix using molecular dynamics simulation, and observed that it exhibited robust stability during simulation frames (Figure 4D). In addition, we also observed that this stretch of βarr2 carboxyl-terminus has a propensity to adopt α-helical conformation even in isolated form i.e., without βarr2 core being present. Interestingly, we did not observe this α-helical structure in D6Rpp-bound βarr1 although the corresponding segment is not resolved in the structure.

**Fig. 4.**
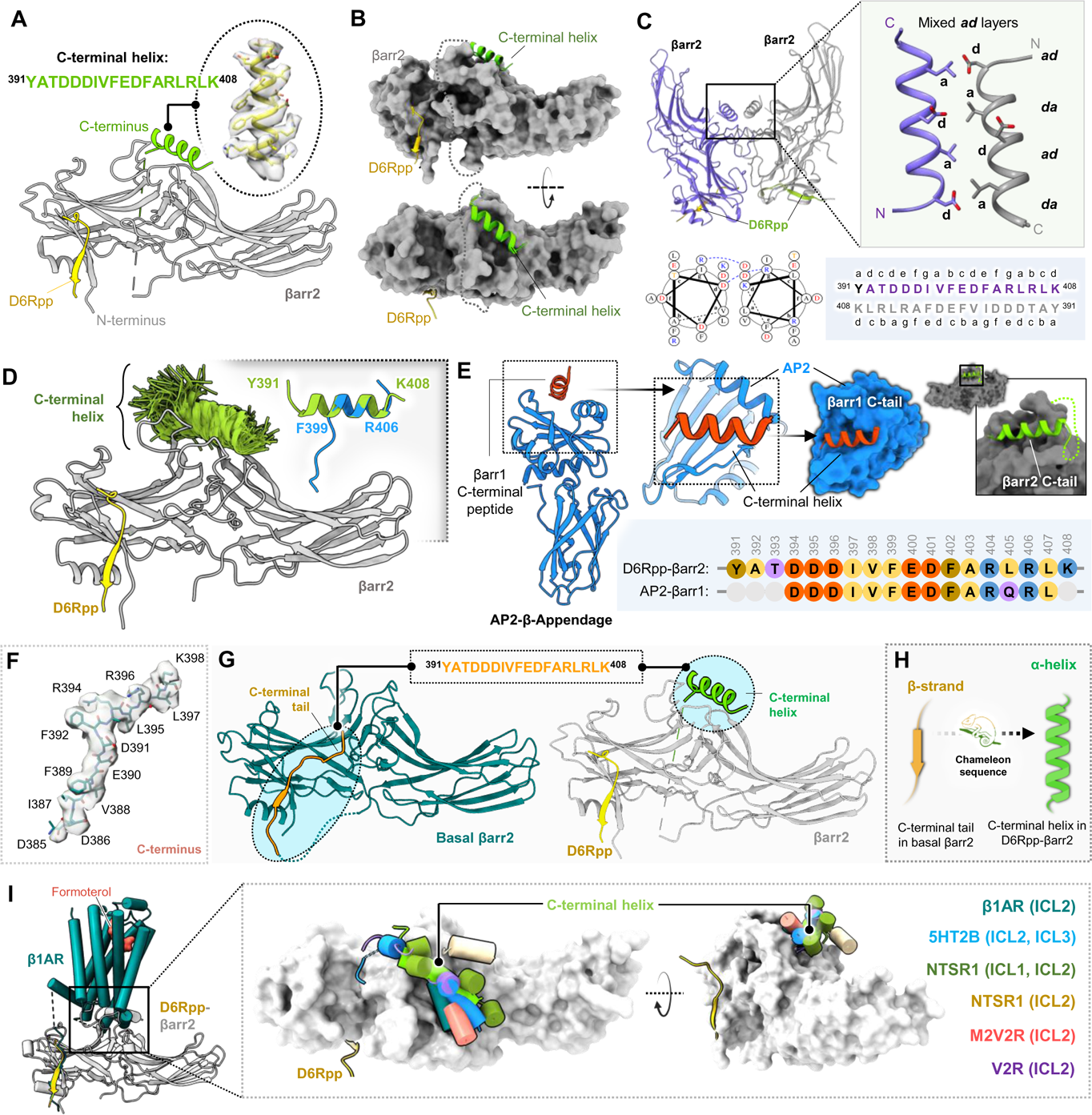
Discovery of a C-terminal helix in D6R activated βarr2. **(A)** Cartoon representation of βarr2 bound to D6R phosphopeptide. βarr2 and D6Rpp are presented in gray and yellow respectively, while the sequence of the C-terminal helix has been provided in an inset. **(B)** D6Rpp-βarr2 structure has been displayed in surface representation in two different views to highlight the pose of the helix. The C-terminal helix (green) and D6Rpp (yellow) are shown as ribbon diagrams. **(C)** Dimeric organization of D6Rpp-βarr2 structure shown in ribbon representation (top left). Formation of anti-parallel coiled-coil (obtained using DrawCoil 1.0 by the C-terminal helix of βarr2 at the dimeric interface (top right) shown as cartoon representation. The anti-parallel coiled-coil exhibits mixed *ad* layers. Helical wheel representation of the anti-parallel coiled-coil shows Asp at position *d* of one helix which forms salt bridge with Arg at position *g* in the other helix (bottom left). Heptad helical representation of the anti-parallel coiled-coil residues in the βarr2 sequence (bottom right). **(D)** MD simulations confirm stability of the distal C-terminal helix/βarr2 interface. Structural snapshots (1 snapshot every 10ns, 7 x 250ns of simulation time) presented here are of the position of the C-tail during simulation. For each residue, frames where it assembles a ⍺-helical conformation are colored green. Fragments of the C-terminal helix can spontaneously assemble a ⍺-helical conformation (right corner, blue cartoon) in 3 out of 4 independent MD simulations (each 2µs) which is overlayed with the crystallized C-tail for comparison (green cartoon). The For each residue, frames where it assembles a helical conformation are colored green. Comparison of a spontaneously assembled helical conformation of the βarr2 C-tail (blue) with that present in the structure (gray). **(E)** Structure of AP2-β-appendage protein in complex with βarr1 C-terminal peptide (PDB 2IV8) has been shown as cartoon representation (left). The βarr1 C-terminal peptide can be seen to adopt similar helical conformation as the C-terminal helix in the D6Rpp bound βarr2 structure (right). The sequence alignment of the C-terminal stretches of βarr1 and βarr2 are shown in inset. **(F)** Cryo-EM density map of the isolated C-terminus of βarr2 has been illustrated. **(G)** The peptide stretch sequence (top) of C-tail in basal βarr2 transforms into a helical conformation in D6Rpp bound state (highlighted in cyan circles). **(H)** The C-tail of βarr2 exhibits a chameleon like property adopting a helical conformation in active state from a β-strand in the basal state. **(I)** Ribbon representation of β1AR-βarr1 structure superimposed with D6Rpp-βarr2 on βarrs (left) shows positioning of C-terminal helix on the central crest of βarrs. Upon structural superimposition with all reported GPCR-βarr1 structures, ICL1/2/3 of various receptors reside on the central crest as C-terminal helix on D6Rpp-βarr2 (right).

It is important to note that in previous structures of activated βarrs, either in complex with phosphopeptides or full-length receptors, either truncated βarrs have been used, or the carboxyl-terminus is not resolved structurally. Even in the crystal structure of βarr2 in its basal conformation, which is used as the only reference for basal conformation in the field, only a part of the carboxyl-terminus is structurally resolved (*40, 41*). Therefore, we also determined the cryo-EM structure of wild-type, full length βarr2, and a significantly longer stretch of the carboxyl-terminus was resolved compared to the previously available crystal structure (Figure 4F-G and Figure S16). Still however, the same stretch of βarr2 adopts a β-strand in its basal conformation, which docks to the N-domain and maintains βarrs in an inactive conformation. Interestingly, a previous structure of β-appendage domain of Adaptin (AP2) in complex with a peptide corresponding to the C-terminus of βarr1 also exhibits an α-helical conformation of the peptide that is positioned onto a groove in the platform sub-domain of β-appendage (Figure 4E) (*42*). Thus, the propensity of the carboxyl-terminus in βarr1 and 2 to adopt α-helical conformation should be explored further.

## Discussion

We note that a cryo-EM structure of a chimeric M2R with engineered V2R carboxyl-terminus (M2-V2R) with βarr1 has been determined previously (*11*), however, the ICL3 of M2R was not resolved in the structure. Therefore, it remains unknown how precisely M2R or other similar GPCRs with short carboxyl-terminus but relatively longer ICL3 engage βarrs (*43*). Our structure of M2R-βarr1 and M2Rpp-βarr2 underscore that the key interaction interface and the activation mechanism remains rather conserved despite distinct domains on the receptor being used to engage βarrs. This essentially starts to provide a structural basis of long-standing questions in the field about how two isoforms of βarrs are able to interact with, and regulate, a broad set of receptors with structurally conserved interface and activation mechanism. We also note from the C3aRpp-βarr1 structure and the comparison of all other structures determined so far of βarr1 and βarr2 pairs bound to the same receptor, underlines a significantly higher inter-domain rotation in βarr2 compared to βarr1 (Figure S17). It is tempting to speculate that this observation provides a molecular mechanism of how class B GPCRs classified based on relatively stable βarr interaction, exhibit apparently higher affinity for βarr2 over βarr1 that was reported almost two decades ago (*44*). Moreover, a direct comparison of M2R-bound βarr1 structure presented here with previously reported M2R-V2R-βarr1 complex reveals the hanging conformation in our complex in terms of βarr1 positioning with respect to the receptor component (Figure S18). This observation further underlines the occurrence of hanging conformation as a major population in the context of native M2R-βarr interaction, and offers a structural framework to design guided experiments in order to probe functional outcomes in future studies. However, the active conformations of βarr1 were similar in terms of the interacting residues on N-domain, key loops, and C-domain rotation values (Figure S19).

The observation of an α-helical conformation in βarr2 upon activation by D6Rpp is intriguing from multiple perspectives. For example, the same conformation is not observed in βarr1, and while this may simply be due to higher flexibility of the carboxyl-terminus in βarr1, it would be anticipated that extensive interactions would allow structural visualization of α-helix if it was being formed. It is intriguing to note that D6Rpp-bound βarr2 exhibits a dimeric assembly while all the previously determined active-like structures such as those bound to V2Rpp, C5aR1pp, M2Rpp, and IP6 reveal a trimeric state (Figure S20 and Table S4). In addition, the α-helix observed in the carboxyl-terminus of βarr2 in D6Rpp-bound state is also absent from the previously determined βarr2 structures. While it cannot be completely ruled out that these differences may arise due to a preferential orientation of the samples on cryo-EM grids, it is tempting to speculate that these differences underscore the conformational signatures in βarrs upon their interaction with GPCRs vs. ACRs, which should be investigated further in subsequent studies. The α-helix in D6Rpp-βarr2 also underscores the “chameleon” nature of the distal carboxyl-terminus to adopt a β-strand in the basal state while transitioning to α-helix upon activation (Figure 4H). Interestingly, such secondary structure switching is also observed for several other proteins that exhibit functional diversity (*45*). It is tempting to speculate that the positioning of α-helix in the central crest of βarr2 may potentially interfere with the core interaction of βarr2 with the receptor although it remains to be experimentally visualized in future studies. This notion is supported by the overlay of D6Rpp-bound βarr2 with previously determined GPCR-βarr structures where either of the ICLs of the receptors appears to clash with the α-helix in βarr2 (Figure 4I and Figure S21). Whether this is a general feature of ACR-βarr interaction or specific to D6R, remains to be examined experimentally in future, possibly through additional structural snapshots and experiments focused to probe conformational dynamics in solution.

We also note that there are several key questions that remain to be answered in the context of GPCR-βarr interaction. For example, there are several prototypical GPCRs that are likely to engage βarrs through their ICL3 but lack P-X-P-P motif, and even some of the ACRs such as CXCR7 and C5aR2 lack this motif in their carboxyl-terminus but they still recruit βarrs. It is also noteworthy that the structural snapshots presented here involve isolated phosphopeptides with defined phosphorylation patterns without the transmembrane core of the receptors. Thus, it is likely that there exist additional mechanisms and/or conformations of βarrs induced by such receptors that remain to be visualize in future studies. As the interaction of receptor core imparts additional conformational changes in βarrs (*46, 47*), it is plausible that the full complexes of receptors and βarrs may exhibit additional conformational changes in βarrs, especially in terms of the positioning of the proximal region of the phosphorylated segment. However, the conserved principle of “P-X-P-P key” to open the “K-K-R-K-R-K lock” is likely to be maintained and guide βarr activation even in the context of full receptors (Figure S22).

In summary, we present novel structural insights into agonist-induced βarr interaction and activation by selected 7TMRs through previously uncharacterized domains namely ICL3, and identify a structural transition in βarr2 carboxyl-terminus from β-strand to α-helix (Figure 5). Taken together, our findings provide important missing information about the current understanding of 7TMR-βarr interaction and signaling with broad implications for GPCR activation, signaling and regulatory paradigms.

**Fig. 5.**
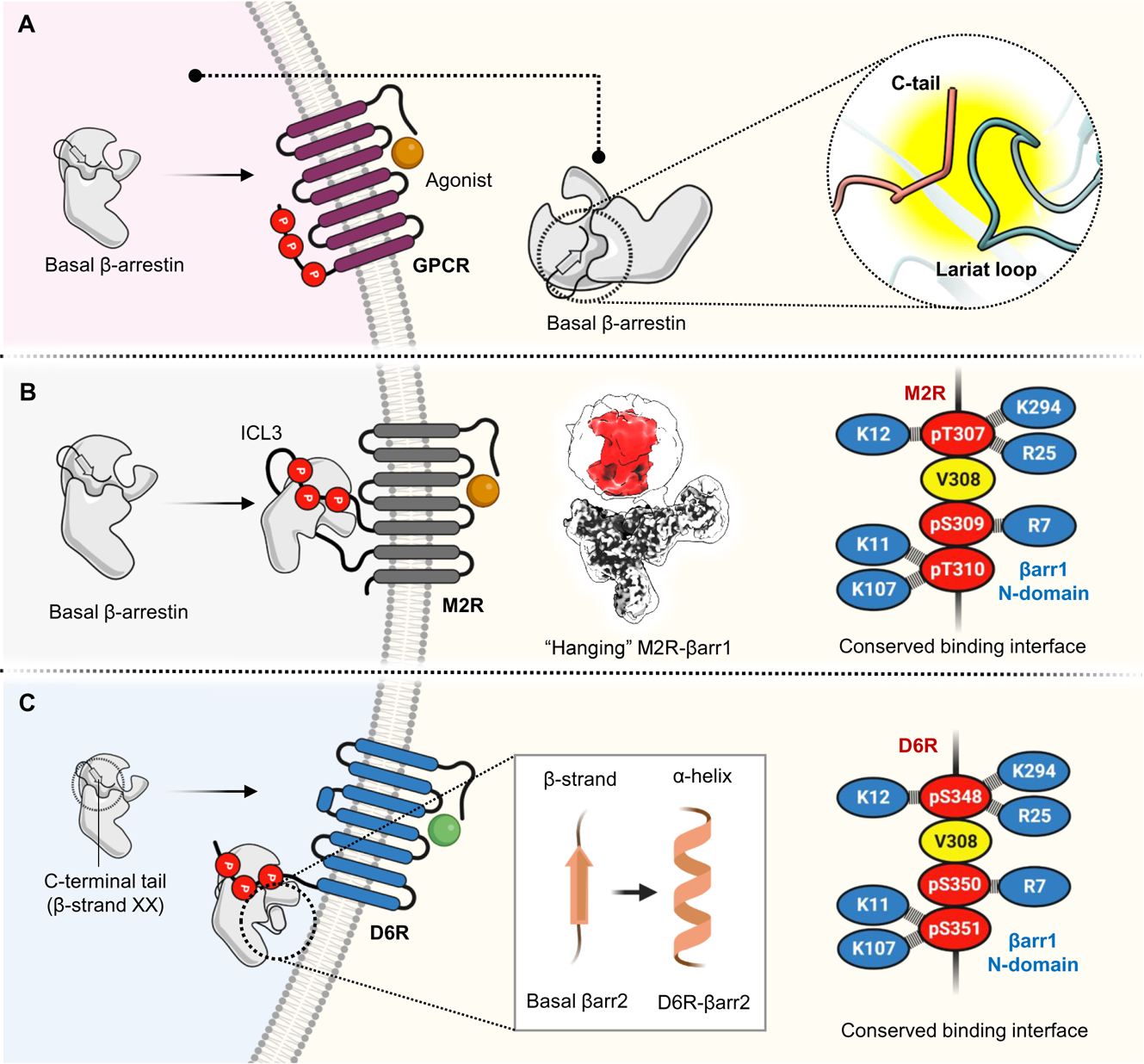
Non-canonical nature of βarr interaction with 7TMRs. **(A)** βarrs in basal state get recruited to phosphorylated GPCRs. The basal conformation of βarrs is stabilized by extensive interactions between the C-terminus and the lariat loop of βarrs. (**B)** M2R-βarr1 adopts a “partially engaged” or “hanging” mode of complex in solution. Despite harboring phosphorylation patterns on long ICL3 in M2R, βarrs engage in similar interaction as in a prototypical receptor. **(C)** βarrs provide a similar set of interacting residues for engaging with the phosphorylated tail of D6R as canonical GPCRs. The C-tail of βarr2 adopts a β-strand conformation in its basal state, whereas it attains an α-helical form upon binding to D6Rpp. Moreover, positioning of the C-terminal helix might sterically clash with ICLs of D6R and could prevent a core-engaged conformation of βarr2 when present in an intact D6R bound complex.

## ACKNOWLEDGMENTS

Research in A.K.S.’s laboratory is supported by the Senior Fellowship of the DBT Wellcome Trust India Alliance (IA/S/20/1/504916) awarded to A.K.S., Science and Engineering Research Board (SPR/2020/000408 and IPA/2020/000405), Council of Scientific and Industrial Research [37(1730)/19/EMR-II], Indian Council of Medical research (F.NO.52/15/2020/BIO/BMS), Young Scientist Award from Lady Tata Memorial Trust, and IIT Kanpur. This work was supported by grants from the JSPS KAKENHI, grant numbers 21H05037 (O.N.), 22K19371 and 22H02751 (W.S.), and 23KJ0491 (F.S.); The Kao Foundation for Arts and Sciences (W.S.); The Takeda Science Foundation (W.S.); The Lotte Foundation (W.S.); and the Platform Project for Supporting Drug Discovery and Life Science Research (Basis for Supporting Innovative Drug Discovery and Life Science Research (BINDS)) from the Japan Agency for Medical Research and Development (AMED), grant numbers JP22ama121012 (O.N.) and JP22ama121002 (support number 3272, O.N.). HDX-MS work in K.Y.C.’s laboratory was supported by grants from the National Research Foundation of Korea funded by the Korean government (NRF-2021R1A2C3003518 and NRF-2019R1A5A2027340). We thank Manisankar Ganguly for assistance with structural analysis, and Sudha Mishra, Annu Dalal and Nashrah Zaidi for help with functional assays. Cryo-EM on basal state βarr2, C3aRpp-βarr1 and, D6Rpp-βarr complexes were performed at the BioEM lab of the Biozentrum at the University of Basel, and we thank Carola Alampi and David Kalbermatter for their excellent technical assistance.

## AUTHOR CONTRIBUTIONS

JM and MKY prepared various complexes used here for structural analysis, JM processed the cryo-EM data together with RB and prepared the figures with input from RB; FKS prepared cryo-EM grids for the M2R complexes, collected and analyzed the data under the supervision of WS and ON; PS carried out the site directed mutagenesis and functional assays; LD performed the HDX-MS experiments under the supervision of KYC; TMS carried MD simulation under the supervision of JS; MC and AR contributed in D6Rpp characterization; VS, SS, and GM contributed in protein purification; MoC prepared the grids and collected cryo-EM data on D6Rpp complexes; AKS supervised the overall project and wrote the manuscript with input from all the authors.

## METHODS

### General reagents, plasmids for cellular assay

Most standard reagents were purchased from Sigma Aldrich unless mentioned. Dulbecco′s Modified Eagle′s Medium (DMEM), Phosphate Buffer Saline (PBS), Trypsin-EDTA, Fetal-Bovine Serum (FBS), Hank’s Balanced Salt Solution (HBSS), and Penicillin-Streptomycin solution were purchased from Thermo Fisher Scientific. HEK-293 cells were purchased from ATCC and cultured in 10% (v/v) FBS (Gibco, Cat. No. 10270-106) and 100U mL^-1^ penicillin and 100µg mL^-1^ streptomycin (Gibco, Cat. No. 15140122) supplemented DMEM (Gibco, Cat. No. 12800-017) at 37°C under 5% CO_2_. For β-arrestin recruitment assays, LgBiT/SmBiT-βarr1/2 and Lg-CAAX construct were used and the same as previously described (*67*). For bystander NanoBiT assay, the cDNA coding region of M2R-WT, M2R-AVAA, and M2R-ANAA with a HA signal sequence, a FLAG tag followed by the N-terminal region of M4 receptor (2-23 residues) at the N-terminus was cloned into pcDNA3.1 vector. To study direct βarr recruitment assay, D6R-WT and D6R-Δ338 harboring SmBiT at the carboxyl-terminus were cloned into the pCAGGS vector. For crosslinking coIP, βarr1 cloned into pCMV vector was used. All DNA constructs were verified by sequencing from Macrogen. The small molecule compound Carbachol was synthesized from Cayman Lifesciences, and CCL7 was purified in the laboratory.

### Expression and purification of βarrs

For expression and purification of βarrs, a previously reported protocol was followed (*48*). In brief, cDNAs of rat βarr1, βarr2^WT^ and bovine βarr2^DM^ (full-length) were cloned into pGEX4T3 vector with GST tag and thrombin cleavage site. An isolated *E. coli* BL21 colony was inoculated into a primary culture of 50mL TB medium supplemented with 100µg mL^-1^ ampicillin. After growing up to a cell optical density at 600nm (OD600) of 0.8-1.0, a secondary culture of 1.5L Terrific Broth media was inoculated from the primary culture and grown till an optical density at 600nm (OD600) of 0.8-1.0. The expression of βarrs were enhanced with 25µM concentration of IPTG and further incubated till 16h at 18°C. Cultures were harvested and stored at -80°C until further use.

Cell lysis was carried out by sonicating the pellets resuspended in lysis buffer 25mM Tris, pH 8.5, 150mM NaCl, 1mM PMSF (phenylmethylsulfonyl fluoride), 2mM Benzamidine, 1mM EDTA (Ethylenediaminetetraacetic acid), 5% Glycerol, 2mM Dithiothreitol (DTT) and 1mg mL^-1^ Lysozyme. The lysate was further spun at 18,000-20,000rpm at 4°C followed by filtration with 0.45µm pore size filter to obtain a clear supernatant. Batch binding was performed overnight with Glutathione resin (Glutathione SepharoseTM 4 Fast Flow, GE Healthcare Cat. no. 17-5132-02) at 4°C. Subsequently, beads bound with proteins were rigorously washed with (25mM Tris, pH 8.5, 150mM NaCl, 2mM DTT and 0.02% n-dodecyl-β-D-maltopyranoside [DDM]) buffer after transferring into Econo columns (Biorad, Cat. no. 7372512). Thrombin at concentration 1unit µL^-1^ was added to the resin slurry at 1:1 (dry resin:cleavage buffer) with cleavage buffer 25mM Tris, pH 8.5, 350mM NaCl and 0.02% DDM, and incubated for 2h at room temperature for on-column cleavage. Pure, tag-free βarrs were eluted and further purified on HiLoad 16/600 Superdex gel-filtration column in running buffer, 25mM Tris, pH 8.5, 350mM NaCl, 2mM DTT and 0.02% DDM. Fractions corresponding to βarrs were pooled and stored at -80°C by adding 10% glycerol until use.

### Expression and purification of Fabs

For expression and purification of Fabs a similar procedure was followed as reported previously (*49*). Briefly, *E. coli* M55244 cells (ATCC) transformed with Fab plasmid were grown in 5mL 2XYT media for overnight 30°C as seed culture. 1L of 2XYT media was further inoculated using 5% of the seed culture and incubated for 8h at 30°C. Post incubation, cells were harvested and resuspended in 1L of CRAP medium already supplied with 100µg mL^-1^ ampicillin, and further incubated for 16h at 30°C. Cells were harvested and subjected to lysis using sonication with buffer, 50mM HEPES, pH 8.0, 500mM NaCl, 0.5% (v/v) Triton X-100, 0.5mM MgCl_2_. The lysate was heated at 65°C in a water bath for 30min and immediately chilled on ice for 5min. To obtain a clear supernatant, lysate was centrifuged for 30min at 20,000g and loaded into a column packed with Protein L resins at room temperature. Post bead binding, washing was performed with 50mM HEPES, pH 8.0, 500mM NaCl buffer. Proteins were eluted with 100mM acetic acid in tubes filled with 1M HEPES, pH 8.0 at 10% of column volume for quick neutralization of eluted proteins. Protein solution was then buffer-exchanged into buffer, 20mM HEPES, pH 8.0, 100mM NaCl using pre-packed de-salting columns (GE Healthcare Cat. no. 17085101). Fabs were then stored at -80°C by adding 10% glycerol until further use.

### Co-immunoprecipitation assay using purified proteins

β-arrestin interaction with phosphopeptides derived from receptors was studied by Co-immunoprecipitation assay. In brief, 5µg of β-arrestin was activated by incubating it with 10 molar excess of individual phosphopeptides on ice for 40min followed by adding 2.5µg of Fab30. The reaction was incubated at room temperature with constant mixing on a tumbler (5rpm) for 1h. 25µL of Protein L beads (Cat. no. Capto™ L resin, GE Healthcare Cat. no. 17547802), pre-equilibrated with binding buffer (20mM HEPES, PH 7.4, 150mM NaCl and 0.01% MNG) was added to each reaction and further incubated for 1h. After 1h, beads were extensively washed with binding buffer and eluted in 30µL 2X SDS dye. 20µL sample was then analyzed on 12% SDS-PAGE, and the intensity of the protein band was quantified by ImageJ (*50*) for statistical analysis. The data were normalized with respect to their respective experimental control and appropriate statistical analyses were performed as indicated in the corresponding figure legend.

### Limited trypsin proteolysis

To quantify the conformational changes in β-arrestin upon binding with differently phosphorylated phosphopeptides derived from the C-terminus of D6R, we performed limited trypsin proteolysis following previously established protocols (*51–53*). Briefly, 10µg of β-arrestin was incubated with a 50-fold molar excess of phosphopeptide for 40min on ice. Activated β-arrestin was digested with TPCK-treated trypsin (Sigma Aldrich, Cat no. T1426) in a 1:100 (trypsin: arrestin) ratio (w/w) for 5-10min at 37°C. The reaction was stopped by transferring 20µL of the reaction mix to another microcentrifuge tube containing 5µL of 5X SDS-protein loading dye. Digestion reactions were analyzed by SDS-PAGE, and digested products were quantified using the Image J program. Trypsin untreated and apo β-arrestin were also taken as controls for every set of experiments.

### Reconstitution of receptor-βarr-Fab complexes from *Sf9* cells

Wild type, full-length, human receptors (M2R, D6R, C3aR) were used for complex reconstitution with βarrs and Fabs. A similar protocol was followed for purification of all the receptor-βarr-Fab complexes. For expression of receptors, the constructs contain haemagglutinin (HA) sequence and FLAG tag followed by a portion of M4R (Muscarinic receptor 4; ANFTPVNGSSGNQSVRLVTSSS), and a 3C protease cleavage site in the N-terminus. Baculoviruses were generated for each receptor till passage P3 stage. For the reconstitution of complexes, receptors were expressed and purified from *Sf9* cells while the other components (βarr and Fab) were added after FLAG elution of receptors. In some cases, viruses were also prepared for βarrs and GRK2/GRK6 till P3 passage. 600mL of *Sf9* cells at 1.8-2.0×10^6^ mL^-1^ density were infected with 12-14mL of receptor, 4-6mL of βarr and 3-5mL of GRK viruses and incubated for 72h at 27°C. The morphology of infected cells was routinely checked under microscope. Cells were stimulated with agonists 1h prior to harvesting. Carbachol (1mM) (Cat. no. 51-83-2, Cayman chemical), CCL7 (1µM, in-house purified) and C3a (1µM, in-house purified) were supplemented to M2R, D6R and C3aR cultures, respectively. Pellets were stored at -80°C until purification.

Before proceeding with purification, expression status for all complex components were checked using western blot analysis. Similar purification steps were followed for all receptor-βarr complexes. Co-expressed culture pellets were resuspended in buffer 20mM HEPES, pH 7.4, 150mM NaCl, 1mM PMSF and 2mM Benzamidine and homogenized using glass dounce-homogenizer for 60 strokes. Fabs were supplemented at a 1.5 molar excess of an estimated receptor amount and kept on stirring for 1h at room temperature. Post incubation, 1% LMNG, 0.01% CHS were added to the lysate and further homogenized for 60 strokes and was incubated for solubilization for 2h at 4°C. Lysate was centrifuged for 30min at 18,000-20,000rpm. The supernatant was filtered with 0.45µm pore-size filter before proceeding for bead binding. Clear lysate was passed onto M1-FLAG resin pre-packed into glass Econo columns (Biorad, Cat. no. 7372512) and allowed to gravity-flow at 1-2mL min^-1^. Extensive washing was done by passing a low-salt buffer (20mM HEPES, pH 7.4, 150mM NaCl, 0.01% LMNG, 0.01% CHS and 2mM CaCl_2_) thrice and with a high-salt buffer (20mM HEPES, pH 7.4, 350mM NaCl, 0.01% LMNG) twice, each with 10mL of volume alternatively. FLAG peptide at concentration of 0.25mg mL^-1^ was added to low-salt buffer for gravity flow elution at around 1mL min^-1^. Fractions were further analyzed on SDS-PAGE and concentrated with 100 MWCO concentrators (Vivaspin, Cytiva Cat. no. 28932319) before gel-filtration chromatography. Superose 6 Increase 10/300 GL (Cytiva Cat. no. 29091596) column was used for further purifying the complexes with a running buffer (20mM HEPES, pH 7.4, 100mM NaCl, 0.00075% LMNG, 0.00025% CHS). Elution fractions corresponding to complexes were analyzed on SDS-PAGE and concentrated to 8-10mg mL^-1^ for negative-staining EM and cryo-EM studies. Respective agonists were kept in excess during all steps and buffers of purification.

### Reconstitution of phosphopeptide-βarr-Fab complexes

A previously published protocol was followed for the phosphopeptide-βarr complex reconstitution (*17*). In brief, phosphopeptides at three molar excesses were added to βarrs and incubated for 30min at room temperature for activation. Post incubation, corresponding Fabs were mixed at 1:1.5 ratio (βarr:Fab) and allowed for complex formation for 90min at room temperature. The reconstituted complexes were further purified on Superose 6 Increase 10/300 GL (Cytiva Cat. no. 29091596) gel-filtration column with a running buffer (20mM HEPES, pH 7.4, 100mM NaCl, 0.00075% LMNG, 0.00025% CHS and 2mM DTT) post concentration with 30,000 MWCO concentrators (Vivaspin, Cytiva Cat. no. 28932361). Fractions corresponding to the complex were pooled, concentrated to desired concentration (8-10mg mL^-1^) and used for negative-staining EM and cryo-EM analysis.

### Glutaraldehyde crosslinking of M2R-βarr1 complex

An on-column cross-linking step was performed to stabilize the pre-formed M2R-βarr1-Fab30 complex. A previously reported protocol was followed with modifications (*35*). Here, pre-packed PD-10 desalting columns (GE Healthcare Cat. no. 17085101) were used instead of gel-filtration columns. The below described protocol has been optimized for 250µL of complex solution. 250µL of glutaraldehyde (1% final concentration) was applied to the pre-equilibrated de-salting column in buffer (20mM HEPES, pH 7.4, 100mM NaCl, 0.00075% LMNG, 0.00025% CHS, 1mM Carbachol) and allowed to gravity-flow. Subsequently, 500µL of running buffer was given twice in sequence. The reconstituted complex sample (250µL) was then allowed to pass through the column with gravity-flow. Immediately after loading the complex sample, two rounds of running buffers (500µL each) were passed down the column. After flow-through of ∼2.5mL, elution was carried out with loading the running buffer and fractions were collected in separate tubes filled with 350µL of 1M Tris, pH 8.0 for quenching additional cross-linking of proteins in proximity. Elution fractions were analyzed on SDS-PAGE and proceeded for further rounds of purification with size-exclusion chromatography after concentration with 100 MWCO concentrators (Vivaspin, Cytiva Cat. no. 28932319). After separating cross-linked aggregates with the Superose 6 Increase 10/300 GL (Cytiva Cat. no. 29091596) gel-filtration column, fractions corresponding to complex were further concentrated and sent for EM analysis.

### Negative-staining EM

Negative-staining EM of all samples were performed to assess complex formation, homogeneity and particle quality prior to grid freezing for cryo-electron microscopy. Negative staining and imaging of the samples were performed in accordance with a previously published protocol (*28*). Briefly, 3.5μL of the protein sample were dispensed on glow discharged carbon/formvar coated 300 mesh Cu (PELCO, Ted Pella) grid, allowed to adsorb for 1min and blotted off using a filter paper. Two separate drops of freshly prepared 0.75% (w/v) uranyl formate stain were set and the grid was gently touched onto the first drop of stain, and immediately blotted off using a filter paper. The grid was then touched onto a second drop of stain for 30s, blotted off and left on the bench on a petri plate for air drying. Imaging was done on a FEI Tecnai G2 12 Twin TEM (LaB6) operating at 120kV and equipped with a Gatan CCD camera (4k x 4k) at 30,000x magnification. Micrographs were collected and processed in Relion 3.1.2 (*54–56*). About 10,000 autopicked particles were autopicked with the gaussian picker, extracted, and subjected to reference free 2D classification.

### Cryo-EM sample preparation and data acquisition

3µL of the samples corresponding to M2R-βarr1 or M2Rpp-βarr2 complexes were dispensed onto glow discharged Quantifoil holey carbon grids (Au R1.2/1.3) and plunged frozen in liquid ethane (-181°C) using a Vitrobot MarkIV maintained at 100% humidity and 4°C. Data were collected on a 300kV Titan Krios microscope (G3i, Thermo Fisher Scientific) equipped with a K3 direct electron detector (Gatan) and BioQuantum K3 imaging filter. Movies were acquired in counting mode across a defocus range of -0.6 to -1.6μm at a pixel size of 0.83Å/px using EPU software (Thermo Fisher Scientific) software. Movies were dose fractionated into 48 frames with a dose rate of approximately 50e^-^/Å^2^.

For the D6Rpp-βarr, C3aRpp-βarr and basal βarr2 complexes, 3µL of the samples were dispensed onto glow discharged Quantifoil holey carbon grids (Cu R2/1 or R2/2) using a Leica GP plunger (Leica Microsystems, Austria) maintained at 90% humidity and 10°C, and vitrified in liquid ethane. A 300kV TFS Titan Krios microscope equipped with Gatan K2 summit direct electron detector (Gatan Inc.) was used to film the cryo-electron microscopy images for the D6Rpp-βarr2-Fab30 complex. SerialEM software was used to automatically capture images in counting mode across a defocus range of 0.5-2.5µm, at a nominal magnification of 165,000x and pixel size of 0.82. A total dose of 56 e^-^/Å^2^ was divided among 40 frames of each movie stack. A 200kV TFS Glacios microscope equipped with a Gatan K3 direct electron detector (Gatan Inc.) was used to collect data for the D6Rpp-βarr1-Fab30, C3aRpp-βarr1-Fab30 and basal βarr2-Fab6 complexes. Each movie stack was dose-fractionated into 40 frames with a total accumulate a total dose of ∼50e^-^/Å^2^ and exposure time of 4s.

### Cryo-EM data processing and model building

Movie frames corresponding to M2R-βarr1 or M2Rpp-βarr2 complex datasets were aligned (4×4 patches) and dose-weighted with RELION’s implementation of the MotionCor2 algorithm (*57*). The motion corrected micrographs were imported into cryoSPARC v3.3.1 or 4.0 and contrast transfer function parameters were estimated with Patch CTF (multi).

For the non-crosslinked M2R-βarr1-Fab30 complex dataset, 31,758 motion corrected micrographs with CTF fit better than 4.5Å were curated and selected for further processing in cryoSPARC v3.3.1. 17,218,446 particles were automatically picked using the blob-picker subprogram, extracted with a box size of 416px (fourier cropped to 64px) and subjected to reference free 2D classification. Clean 2D classes containing 4,815,631 particles with conformations corresponding to receptor-βarr complexes were selected and re-extracted with a box size of 416px (fourier cropped to 256px). Subsequent ab-initio reconstruction and heterogeneous refinement yielded a 3D class with 34% of the particles and features of GPCR-βarr hanging conformation. Particle projections from this 3D class were extracted with full box size (416px) and subjected to non-uniform refinement to yield a map with clear density and secondary features corresponding to βarr-Fab30 portion but not very well-defined micellar density, suggesting flexibility in the micelle region of the map. Particle subtraction was performed on the particle projections with mask on the micelle, followed by local refinement with mask on the β-arrestin and variable domain of Fab30. This yielded a locally refined map (voxel size of 0.83Å/px) with an overall resolution of 3.1Å in accordance with the gold standard Fourier Shell Correlation (FSC = 0.143) criteria. DeepEMhancer (*58*) available at the COSMIC cryo-EM webserver was used for map sharpening to improve the interpretability and remove the light directional (resolution) anisotropy exhibited by the final map.

For the crosslinked M2R-βarr1-Fab30 complex dataset, a total of 5,235,492 particles were automatically picked, extracted with a box size of 416px (fourier cropped to 64px), and subjected to 2D classification, ab-initio reconstruction, and heterogeneous refinement. The following steps were the same as those used for the non-crosslinked M2R-βarr1-Fab30 complex dataset. The particle projections corresponding to the best 3D class were re-extracted with a box size of 416px (fourier cropped to 288px). The re-extracted particles were subjected to focused 3D classification (without alignment) with a mask on the βarr-Fab30 component, followed by homogeneous refinement yielding a map with an overall resolution of 3.5Å. The map so obtained was subjected to local refinement with mask on β-arrestin and variable domain of Fab30 portion resulting in a map with an overall resolution of 3.2Å (voxel size of 1.2Å/px) with the gold standard Fourier Shell Correlation using the 0.143 criterion. As for the crosslinked M2R-βarr1-Fab30 complex dataset, the final map exhibited a small degree of anisotropy, which was also corrected through map sharpening with DeepEMhancer.

For the M2Rpp-βarr2-Fab30 complex dataset, 1,861,553 particles were autopicked from 2,596 motion corrected micrographs using blob-picker and extracted with a box size of 416px (fourier cropped to 64px). Reference free 2D classification yielded class averages with clear secondary features corresponding to a trimeric assembly. Selected 2D averages containing 1,861,553 particles were re-extracted with a box size of 416px (fourier cropped to 288px) and subjected to ab-initio reconstruction followed by heterogenous refinement yielding 2 classes. Non-uniform refinement with C3 symmetry followed by local refinement with mask on the β-arrestin and variable domain of Fab30 yielded a map with an overall resolution of 2.9Å (voxel size = 1.2Å/px) according to the FSC = 0.143 criterion.

For the D6Rpp-βarr, C3aRpp-βarr and basal βarr2 complex datasets, all data processing steps were performed in cryoSPARC 3.3.2 or 4.0 unless otherwise stated. Patch motion correction (multi) was used to perform beam-induced motion correction on the dose-fractionated movie stacks, and Patch CTF estimation (multi) was used to estimate the contrast transfer function parameters.

For the D6pp-βarr2-Fab30 dataset, 9,977 dose weighted, motion corrected micrographs with CTF fit resolution better than 4.5Å were chosen for further processing. 496,954 particles were autopicked, extracted with a box size of 480px (fourier cropped to 64px) and subjected to reference-free 2D classification to eliminate junk particles. 337,137 particle projections corresponding to 2D class averages with evident secondary features were subjected to ab-initio reconstruction yielding 3 classes. Following heterogenous refinement, the 3D class with characteristics of a dimeric βarr-Fab30 complex containing 83,459 particles (58% of the total particles) was subjected to nun-uniform refinement with C2 symmetry followed by local refinement with a mask to remove the constant zone of Fab30. This resulted in a coulombic map with a global resolution of 3.2 at 0.143 FSC cut-off.

For the D6pp2-βarr1-Fab30 dataset, 5,300,908 particles were initially picked from the total of 9,698 micrographs using the blob-picker sub-program. These particles were extracted with a box size of 480px (fourier cropped to 64px) and subjected to several rounds of 2D classifications. The best 2D averages containing 511,711 particles were re-extracted with a box size of 480px (fourier cropped to 288px) and subjected to ab-initio reconstruction and heterogenous refinement yielding two models. The 3D class containing a dimeric architecture and defined secondary features (369,871 particles) was selected for non-uniform refinement and successive local refinement with mask on the β-arrestin molecule and Fab30 variable domain. The final local refinement yielded a map with a global resolution of 3.4Å, according to the FSC at 0.143 criterion.

For the C3aRpp-βarr1-Fab30 complex dataset, two independent data collection sessions – 7,246 movies (untilted) and 6,192 movies (45° tilted) were performed to solve the preferred orientation issue which arose during initial data processing. Particles were picked with blob-picker from both datasets independently, extracted with a box size of 480px (fourier cropped to 64px) and subjected to several rounds of 2D classification to eliminate noisy particles. Particles corresponding to the clean classes from both datasets were then selected, merged, and re-extracted with a box size of 480px (fourier cropped to 288px). The re-extracted particles were then used for ab-initio reconstruction and subsequent heterogenous refinement yielding two models. The 3D class with features of a dimeric complex containing 252,613 particles were then subjected to 3D classification without alignment, followed by non-uniform refinement and local refinement with a mask on the dimeric complex with imposed C2 symmetry. The final map (voxel size of 1.46Å/px) exhibited slight directional anisotropy which was corrected through map sharpening using DeepEMhancer.

For the basal state β-arrestin2-Fab6 complex dataset, 7,887,274 particles were autopicked from 12,586 motion corrected micrographs, extracted with a box size of 360px (fourier cropped to 64px) and subjected to several rounds of 2D classification to yield class averages with clear secondary features. The particles corresponding to the best classes were selected and extracted with a box size of 416px (fourier cropped to 256px) for subsequent ab-initio reconstruction and heterogenous refinement. 506,938 particles corresponding to the best 3D class were subjected to non-uniform refinement yielding a map with 3.7Å overall resolution. Subsequent masked local refinement resulted in a final map with an overall resolution of 3.5Å (1.2347Å/px) as estimated by the gold standard fourier shell correlation using 0.143 criterion.

The β-arrestin molecule and Fab30 were masked for local 3D refinement, which resulted in more distinct densities in the pliable areas, including the loops, and facilitated model construction in the coulombic densities. Local resolution estimates of all maps were calculated using the Blocres module of cryoSPARC and their complementary half maps. All maps were sharpened using Phenix’s “Autosharpen maps” (*59, 60*) tool or DeepEMhancer to improve maps for model building. Detailed schematic pipeline for data processing have been included as Figures S3-S9.

### Model building and refinement

Sharpened maps were used for model building, refinement, validation, and successive structural analysis. Protomeric coordinates of βarr1 were obtained from previously solved cryo-EM structure of C5aR1pp-βarr1-Fab30 complex (PDB 8GO8), while the coordinates of βarr2 and Fab30 were adapted from the cryo-EM structure of V2Rpp-βarr2-Fab30 complex (PDB 8GOC). The initial model of Fab6 was generated in MODELLER (*61*) with the coordinates of Fab30 from 8GOC. These initial models of βarrs and Fabs were docked into the individual coulombic maps with Chimera (*62, 63*), followed by flexible fitting of the docked models with the “all atom refine” module in COOT (*64*). Phosphopeptide residues were built manually. The models obtained were refined with Phenix real-space refinement with imposed secondary structural restraints against the coulombic maps. The final statistics of all models were evaluated with Molprobity (*65*) included within Phenix comprehensive validation job with the final refined models as input. All structural figures used in the manuscript were prepared using either Chimera or ChimeraX (*63*). Data collection, processing and refinement statistics have been included as Table S1.

### Hydrogen/deuterium exchange mass spectrometry (HDX-MS)

Protein samples were prepared at a final concentration of 35-40μM in 20mM HEPES pH 7.4, 150mM NaCl, and 1mM DTT. For phosphopeptide binding, 10-fold excess concentration of D6Rpp was added to β-arrestins and incubated for 30min at room temperature (23-25°C). Hydrogen/deuterium exchange was initiated by mixing 3μL of protein samples with 27μL D_2_O buffer (20mM HEPES pH 7.4, 150mM NaCl, and 10% glycerol in D_2_O) and incubating the mixtures for 10, 100, 1000 and 10000s on ice. At each time point, 30μL of ice-cold quench buffer (60mM NaH_2_PO_4_, pH 2.01, 10% glycerol) was added to quench the deuterium exchange reaction. For non-deuterated samples by mixing 3μL of protein samples with 27μL of H_2_O buffer (20mM HEPES, pH 7.4, 150mM NaCl, and 10% glycerol in H_2_O), followed by quench steps as described above. After injection, the quenched samples were sent for digestion via an immobilized pepsin column (2.1× 30mm) (Life Technologies, Carlsbad, CA, USA) at 100mL min^-1^ with 0.05% formic acid in H_2_O at 12°C. Peptic peptides were transmitted to a C18 VanGuard trap column (1.7μm × 30mm) for desalting with 0.05% formic acid in H_2_O, and then separated by ultra-pressure liquid chromatography through an Acquity UPLC C18 column (1.7μm, 1.0 × 100mm) at 40mL min^-1^ with an acetonitrile gradient of 8-85% B over 8.5min. Mobile phase A was 0.1% formic acid in H_2_O and mobile phase B was 0.1% formic acid in acetonitrile. Buffers were adjusted to pH 2.5 and system was maintained at 0.5°C (except pepsin digestion at 12°C) to minimize the back-exchange of deuterium to hydrogen. Mass spectral analyses were performed by using a Xevo G2 quadrupole time-of-flight (Q-TOF) equipped with a standard ESI source in MS E mode (Waters) in positive ion mode. The capillary, cone, and extraction cone voltages were set at 3kV, 40V, and 4V, respectively. Source and desolvation temperatures were set at 120°C and 350°C, respectively. Trap and transfer collision energies were set to 6V and the trap gas flow rate was set at 0.3mL min^-1^. Sodium iodide (2µg µL^-1^) was used to calibrate the mass spectrometer, and [Glu1]-Fibrinopeptide B (200fg µL^-1^) in MeOH:water (50:50 (v/v)+1% acetic acid) was used for lock-mass correction. The ions at mass-to-charge ratio (m/z) of 785.8427 were monitored at a scan time of 0.1s with a mass window of ±0.5Da. The reference internal calibrant was introduced at a flow rate of 20µL min^-1^, and all spectra were automatically corrected using lock-mass. Two independent interleaved acquisition functions were created. The first function, typically set at 4eV, collected low-energy or unfragmented data, whereas the second function collected high-energy or fragmented data typically obtained using a collision ramp from 30–55eV. Ar gas was used for collision-induced dissociation (CID). Mass spectra were acquired in the range of m/z 100–2000 for 10min. Peptides from non-deuterated samples were identified by ProteinLynx Global Server 2.4 (Waters) with variable methionine oxidation modification and a peptide score of 6. Deuterium uptake levels of each peptide were determined by measuring the centroid of the isotopic distribution via DynamX 3.0 (Waters). All the data was obtained from at least three independent experiments. The summary of HDX-MS profiles and uptake levels of all the analyzed peptides are listed in the Table S5.

### Surface expression assay

Receptor surface expression in respective assays was measured by whole cell-based surface ELISA as previously described (*66*). Briefly, transfected cells were seeded in 0.01% poly-D-Lysine pre-treated 24-well plate at a density of 2×10^5^ cells well^-1^ and incubated for 24h at 37°C. After 24h, growth media was aspirated, and washed once with ice-cold 1XTBS, followed by fixation with 4% PFA (w/v in 1XTBS) on ice for 20min. After fixation, three times washing with 1XTBS (400μL in each wash) followed by blocking with 1% BSA (w/v in 1XTBS) at room temperature for 90min. Afterward, 200μL anti-FLAG M2-HRP was added and incubated for 90min (prepared in 1% BSA, 1:10,000) (Sigma, Cat. no. A8592). Post antibody incubation, three times washed with 1%BSA (prepared in 1XTBS) followed by development of signal by treating cells with 200μL TMB-ELISA (Thermo Scientific, Cat no. 34028) until the light blue color appeared. After that, signal was quenched by transferring the blue-colored solution to a 96-well plate containing 100μL 1M H_2_SO_4_. Absorbance of the signal was measured at 450nm using a multi-mode plate reader. Next, cells were washed two times with 200μL 1XTBS followed by incubation with 0.2% Janus Green (Sigma; Cat no. 201677) w/v for 15min. By washing with distilled water excess stains were removed. After washing, 800μL of 0.5N HCl was added to elute the stain. After elution, 200μL of the solution was transferred to a 96-well plate, and at 595nm, absorbance was recorded. Data were analyzed by calculating the ratio of absorbance at 450/595 followed by normalizing the value of pcDNA transfected cells reading as 1. Normalized values were plotted using GraphPad Prism v 9.5.0 software.

### NanoBiT-based βarr recruitment assay

Plasma membrane localization of βarr upon stimulation of M2R and D6R with respective agonists were measured by a bystander and direct physical recruitment NanoBiT-based assay, respectively, following previously described protocols (*67, 68*). For M2R βarr recruitment study, HEK-293 cells were transfected with 3µg of above mentioned M2R constructs along with N-terminally SmBiT tagged βarr1/2 constructs (3.5µg), and the plasma membrane localization tag CAAX (5µg) harboring LgBiT at the N-terminus using transfection reagent polyethyleneimine (PEI) linear at DNA:PEI ratio of 1:3. For βarr recruitment study downstream of D6R, HEK-293 cells were cotransfected with D6RWT and truncation constructs harboring SmBiT at the C-terminus and βarr1/2 constructs (3.5µg) with N-terminally fused LgBiT. 0.25µg of D6R^WT^, 3.5µg of D6R^Δ338^ were transfected, to match the cell surface expression level. Post 16-18h of transfection, cells were trypsinized, and resuspended in the NanoBiT assay buffer consisting of 1XHBSS, 0.01% BSA, 5mM HEPES, pH 7.4, and 10μM coelenterazine (GoldBio, Cat. no. CZ05). After resuspension of the pellet, cells were seeded in an opaque flat bottom white 96 well plate at a density of 1×10^5^ cells well^-1^. Next, cells were incubated for 120min (90min at 37°C, followed by 30min at room temperature). Post incubation, basal level luminescence readings were taken, followed by ligand addition. For dose-response assay, ligand concentrations ranging from 100pM to 100μM for carbachol and 1pM to 1μM for CCL7 were prepared in the buffer composed of 1XHBSS, 5mM HEPES, pH 7.4, and cells were stimulated with varying doses of indicated ligands. Luminescence upon stimulation was recorded up to 20 cycles by a multimode plate reader. For analysis, average data from the 5 cycles with the maximum reading is used and normalized with respect to the signal of minimal ligand concentration as 1 and plotted using nonlinear regression analysis in GraphPad Prism v 9.5.0 software.

### Chemical cross-linking and co-immunoprecipitation

Agonist-induced βarr recruitment downstream of M2R^WT^ and mentioned mutants was performed by chemical crosslinking following previously published protocol (*69*). Briefly, HEK-293 cells were co-transfected with N-terminally FLAG tagged receptor and βarr1. After 48 h of transfection, cells were serum starved for another 6h, followed by stimulation with 100μM Carbachol. The cell pellet was resuspended in lysis buffer (20mM HEPES, pH 7.4, 100mM NaCl, 0.1mM PMSF, 0.2mM Benzamidine, and 1X Phosphatase inhibitor cocktail) and then lysed in a homogenizer. For crosslinking, freshly prepared crosslinker DSP (3,3′-Dithiodipropionic acid di(N-hydroxysuccinimide ester) (Sigma, Cat. no. D3669) was used at a concentration of 1.5mM. After adding DSP, the sample was incubated for 40 min at room temperature. Post crosslinking, the reaction was quenched by adding 1mM Tris, pH 8.0, and 1% MNG (maltose neopentyl glycol) was added for solubilization for 1h. The bait for this coIP was FLAG M1 antibody coupled beads; beads were pre-equilibrated with buffer consisting of 20mM HEPES, pH 7.4, and 150mM NaCl. After solubilization, spin the lysate at 15,000rpm for 15 min. The supernatant was loaded in the beads for binding, followed by washing, and finally eluted using FLAG-EDTA buffer (20mM HEPES, 150mM NaCl, 2mM EDTA, 0.01% MNG, 250 mg mL^-1^ FLAG peptide). After adding elution buffer, incubate for 30min and flick gently at 10min intervals. The signal was probed by using immunoblotting technique. To probe βarr, βarr1/2 monoclonal anti-rabbit antibody (1:5000, CST, Cat. no. 4674) was used. Blots were stripped, and probe the receptor using anti-FLAG peroxidase coupled antibody (1:2000, Sigma, Cat. no. A8592). Data were quantified using ImageLab software (Bio-Rad) and analysed by dividing signal for βarr by receptor signal followed by normalizing 30min signal for M2RWT as 100%. Data was plotted in GraphPad Prism v 9.5.0 software.

### Molecular dynamics simulations

Residue protonation was assigned using Protonate 3D available within the MOE package (www.chemcomp.com). Complexes were solvated with TIP3P waters containing a 0.15 concentration of NaCl ions. System parameters were derived from Charmm36M (*70*) and subsequent simulations were run using the ACEMD3 engine (*71*). Each system underwent an initial 20ns equilibration run in conditions of constant pressure and temperature (pressure kept constant at 1.01325 bar with the Berendsen barostat), with a timestep of 2fs and restraint applied to protein backbone atoms. Temperature was maintained constant at 310K using the Langevin thermostat, hydrogen bonds were restrained using the RATTLE algorithm. Non-bonded interactions were cut-off at 9Å, with a smooth switching function applied at 7.5Å. To simulate the stability of the βarr2/C-tail interface as well as stability of the dimer with and without the C-tail we have utilized the D6Rpp-βarr2 structure obtained within this study as a starting point. To simulate βarr2 folding we have started with the fully unfolded βarr2 C-tail fragment (residues 392 to 408). To simulate the interface between adaptin and the βarr2 C-tail we have utilized the deposited structure (PDB 2IV8) as a starting point.

