## Supplemental Figures for "Molecular insights into atypical modes of β-arrestin interaction with seven transmembrane receptors"

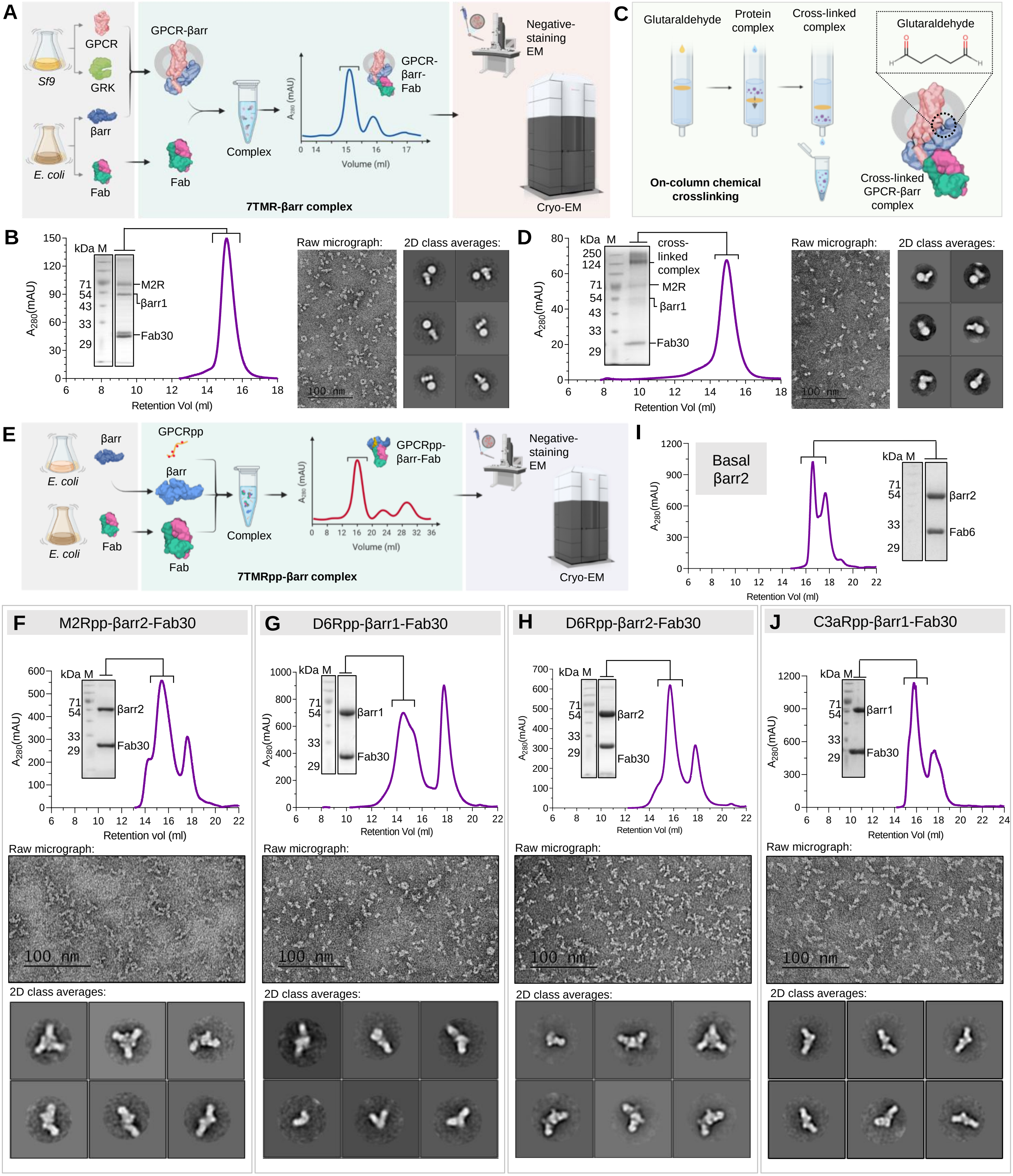
A
Sf9
GPCR
GRK
βarr
E. coli
Fab
GPCR-βarr
Complex
GPCR-βarr-Fab
Volume (ml)
Negative-staining EM
Cryo-EM
7TMR-βarr complex
C
Glutaraldehyde
Protein complex
Cross-linked complex
On-column chemical crosslinking
Cross-linked GPCR-βarr complex
B
M2R
βarr1
Fab30
Retention Vol (ml)
Raw micrograph:
2D class averages:
D
cross-linked complex
E
GPCRpp
GPCRpp-βarr-Fab
7TMRpp-βarr complex
I
Basal βarr2
Fab6
F
M2Rpp-βarr2-Fab30
G
D6Rpp-βarr1-Fab30
H
D6Rpp-βarr2-Fab30
J
C3aRpp-βarr1-Fab30
Retention vol (ml)
100 nm

**Figure S1. Reconstitution of βarr complexes.**

**(A)** Schematic outline of receptor-βarr complex expression and purification protocol. **(B)** Gel-filtration profile, SDS-PAGE analysis (in inset) and visualization through negative-staining EM (representative micrograph to the left, selected 2D class averages ~~in~~ to the right). **(C)** An outline of on-column cross-linking of M2R-βarr1 complex. (D) Gel-filtration profile, SDS-PAGE analysis (in inset) and visualization through negative-staining EM (representative micrograph to the left, selected 2D class averages to the right). **(E)** Schematic outline of in-vitro reconstitution of phosphopeptide-βarr complexes. Gel-filtration profiles followed by SDS-PAGE analysis of **(F)** M2Rpp-βarr2-Fab30 complex, **(G)** D6Rpp-βarr1-Fab30 complex, **(H)** D6Rpp-βarr2-Fab30 complex, **(I)** βarr2-Fab6 complex and **(J)** C3aRpp-βarr1-Fab30 complex, (Scale bar: 100nm).

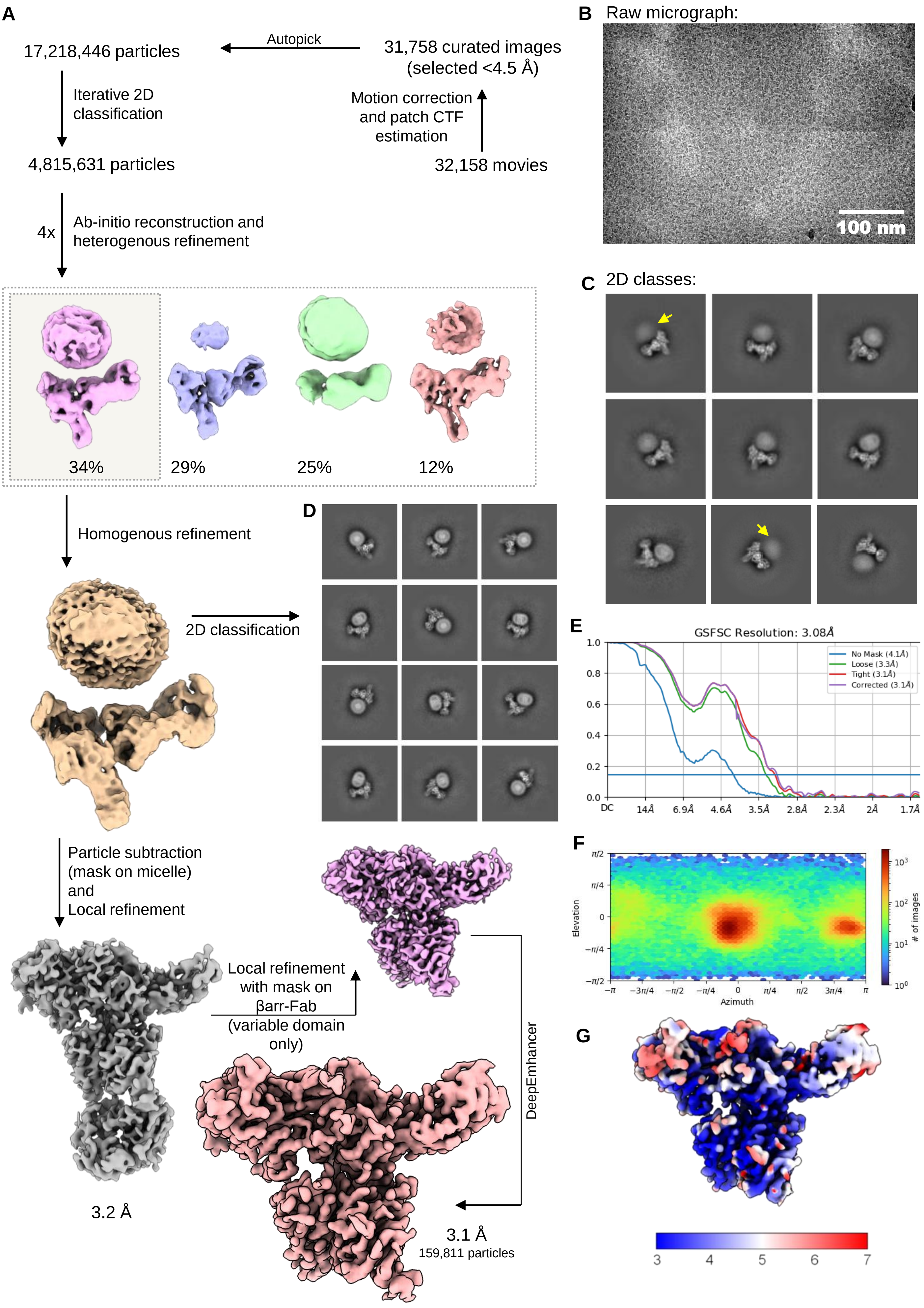
A
17,218,446 particles
Autopick
31,758 curated images (selected <4.5 Å)
Iterative 2D classification
Motion correction and patch CTF estimation
4,815,631 particles
32,158 movies
4x
Ab-initio reconstruction and heterogenous refinement
34%
29%
25%
12%
Homogenous refinement
2D classification
Particle subtraction (mask on micelle) and Local refinement
Local refinement with mask on βarr-Fab (variable domain only)
DeepEmhancer
3.2 Å
3.1 Å
159,811 particles
B
Raw micrograph:
100 nm
C
2D classes:
D
E
GSFSC Resolution: 3.08Å
No Mask (4.1Å)
Loose (3.3Å)
Tight (3.1Å)
Corrected (3.1Å)
DC
14Å
6.9Å
4.6Å
3.5Å
2.8Å
2.3Å
2Å
1.7Å
F
Elevation
Azimuth
### of images
G
3
4
5
6
7

**Figure S2. Cryo-EM reconstruction of the M2R-βarr1-Fab30 (non-cross linked) complex.**

**(A)** Schematic representation of cryo-EM data processing pipeline. **(B)** Representative motion corrected micrograph. **(C)** Selected 2D class averages with flexible micelle density highlighted in yellow arrows. **(D)** Selected 2D class averages showing rigid detergent micelle densities. **(E)** Gold standard fourier shell correlation curve (GFSC) at 0.143 cut-off was used to determine the overall resolution of the map. **(F)** Angular plot of the particles used for 3D reconstruction. **(G)** Local resolution map of the 3D reconstruction in front view. (Scale bar: 100nm).

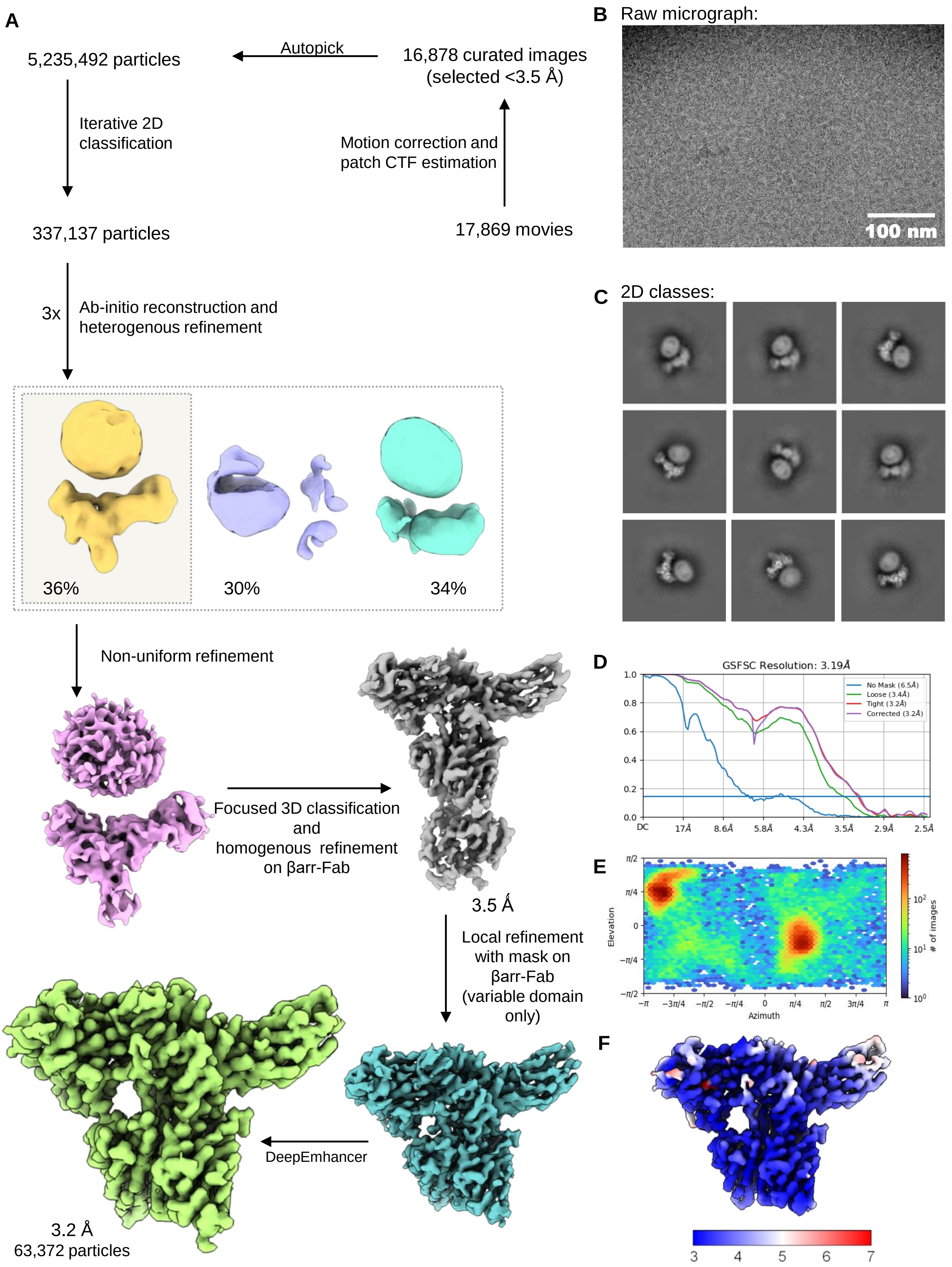

A
5,235,492 particles
Autopick
16,878 curated images
(selected <3.5 Å)
Iterative 2D classification
Motion correction and patch CTF estimation
337,137 particles
17,869 movies
3x
Ab-initio reconstruction and heterogenous refinement
36%
30%
34%
Non-uniform refinement
Focused 3D classification and homogenous refinement on βarr-Fab
3.5 Å
Local refinement with mask on βarr-Fab (variable domain only)
DeepEmhancer
3.2 Å
63,372 particles
B
Raw micrograph:
100 nm
C
2D classes:
D
GSFSC Resolution: 3.19Å
No Mask (6.5Å)
Loose (3.4Å)
Tight (3.2Å)
Corrected (3.2Å)
E
Elevation
Azimuth
### of images
F
3
4
5
6
7

**Figure S3. Cryo-EM reconstruction of the M2R-βarr1-Fab30 (cross linked) complex.**

**(A)** Schematic representation of cryo-EM data processing pipeline. **(B)** Representative motion corrected micrograph. **(C)** Selected 2D class averages. **(D)** Gold standard fourier shell correlation curve (GFSC) at a threshold of 0.143 was used to determine the overall resolution of the map. **(E)** Angular plot of the particles used for 3D reconstruction. **(F)** Local resolution map of the 3D reconstruction in front view. (Scale bar: 100nm).

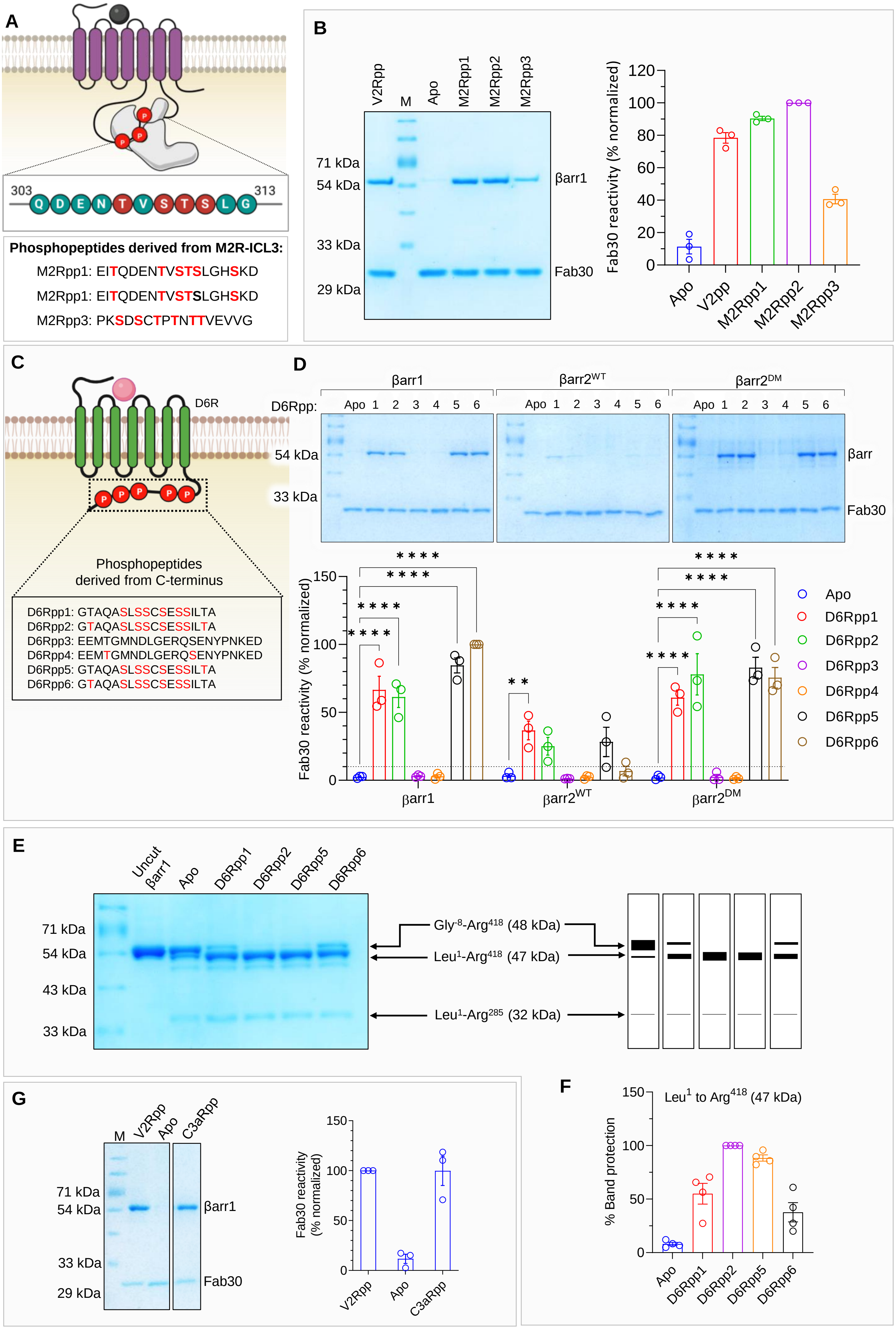
A
303
313
Q D E N T V S T S L G
Phosphopeptides derived from M2R-ICL3:
M2Rpp1: EITQDENTVSTSLGHSKD
M2Rpp1: EITQDENTVSTSLGHSKD
M2Rpp3: PKSDSCTPTNTTVEVVG
B
V2Rpp
M
Apo
M2Rpp1
M2Rpp2
M2Rpp3
71 kDa
54 kDa
33 kDa
29 kDa
βarr1
Fab30
Fab30 reactivity (% normalized)
Apo
V2pp
M2Rpp1
M2Rpp2
M2Rpp3
C
D6R
Phosphopeptides derived from C-terminus
D6Rpp1: GTAQASLSSCSESSILTA
D6Rpp2: GTAQASLSSCSESSILTA
D6Rpp3: EEMTGMNDLGERQSENYPNKED
D6Rpp4: EEMTGMNDLGERQSENYPNKED
D6Rpp5: GTAQASLSSCSESSILTA
D6Rpp6: GTAQASLSSCSESSILTA
D
βarr1
βarr2WT
βarr2DM
D6Rpp:
Apo 1 2 3 4 5 6
54 kDa
33 kDa
βarr
Fab30
Fab30 reactivity (% normalized)
Apo
D6Rpp1
D6Rpp2
D6Rpp3
D6Rpp4
D6Rpp5
D6Rpp6
E
Uncut βarr1
Apo
D6Rpp1
D6Rpp2
D6Rpp5
D6Rpp6
71 kDa
54 kDa
43 kDa
33 kDa
Gly-8-Arg418 (48 kDa)
Leu1-Arg418 (47 kDa)
Leu1-Arg285 (32 kDa)
F
Leu1 to Arg418 (47 kDa)
% Band protection
Apo
D6Rpp1
D6Rpp2
D6Rpp5
D6Rpp6
G
M
V2Rpp
Apo
C3aRpp
71 kDa
54 kDa
33 kDa
29 kDa
βarr1
Fab30
Fab30 reactivity (% normalized)
V2Rpp
Apo
C3aRpp

**Figure S4. Phosphopeptide design for driving βarr activation.**

**(A**) Cartoon representation of design of phosphopeptides derived from ICL3 of M2R. The stretch of ICL3 resolved in the cryo-EM structure is shown in an inset box. **(B)** Fab30 reactivity to M2Rpps as assessed though co-IP assay taking as V2Rpp as control (left). Densitometric analyses of the co-IP assay. **(C)** Designed phosphopeptides derived from D6R. **(D)** Co-immunoprecipitation assay to assess the Fab30 reactivity for D6Rpp activated βarr1/βarr2WT/βarr2DM (top). Densitometry-based quantification (mean ± SEM; n=3; normalized with respect to Fab30 reactivity for activated βarr1 activated by D6Rpp6 signal as 100% (lower panel) (2way ANOVA, Tukey's multiple comparisons test). The exact p values are as follows: reactivity for βarr1- Apo vs. D6Rpp1 ($p < 0.0001$), Apo vs. D6Rpp2 ($p < 0.0001$), Apo vs. D6Rpp5 ($p < 0.0001$), Apo vs. D6Rpp6 ($p < 0.0001$), reactivity for βarr2WT-Apo vs. D6Rpp1 ($p = 0.0067$), reactivity for βarr2DM- Apo vs. D6Rpp1 ($p < 0.0001$), Apo vs. D6Rpp2 ($p < 0.0001$), Apo vs. D6Rpp5 ($p < 0.0001$), Apo vs. D6Rpp6 ($p < 0.0001$). (**$p < 0.01$, ****$p < 0.0001$, ns = non-significant). **(E)** Limited trypsin proteolysis of βarr1 was performed to assess the activation potential of phosphopeptides. The presence and absence of peptides are mentioned in the top of SDS-PAGE analysis gel. A representative gel from four independent experiments (left panel) and a schematic of the proteolysis pattern corresponding to 1:25 ratio of trypsin:βarr1 (right panel) is shown here. **(F)** Densitometry-based quantification (mean ± SEM) of indicated bands from four independent experiments, normalized with respect to D6Rpp2 condition (treated as 100%). **(G)** Fab30 reactivity to C3aRpp as assessed though co-IP assay taking as V2Rpp as control (left). Densitometric analysis (mean ± SEM) of corresponding Fab30 reactivity with V2Rpp as 100%.

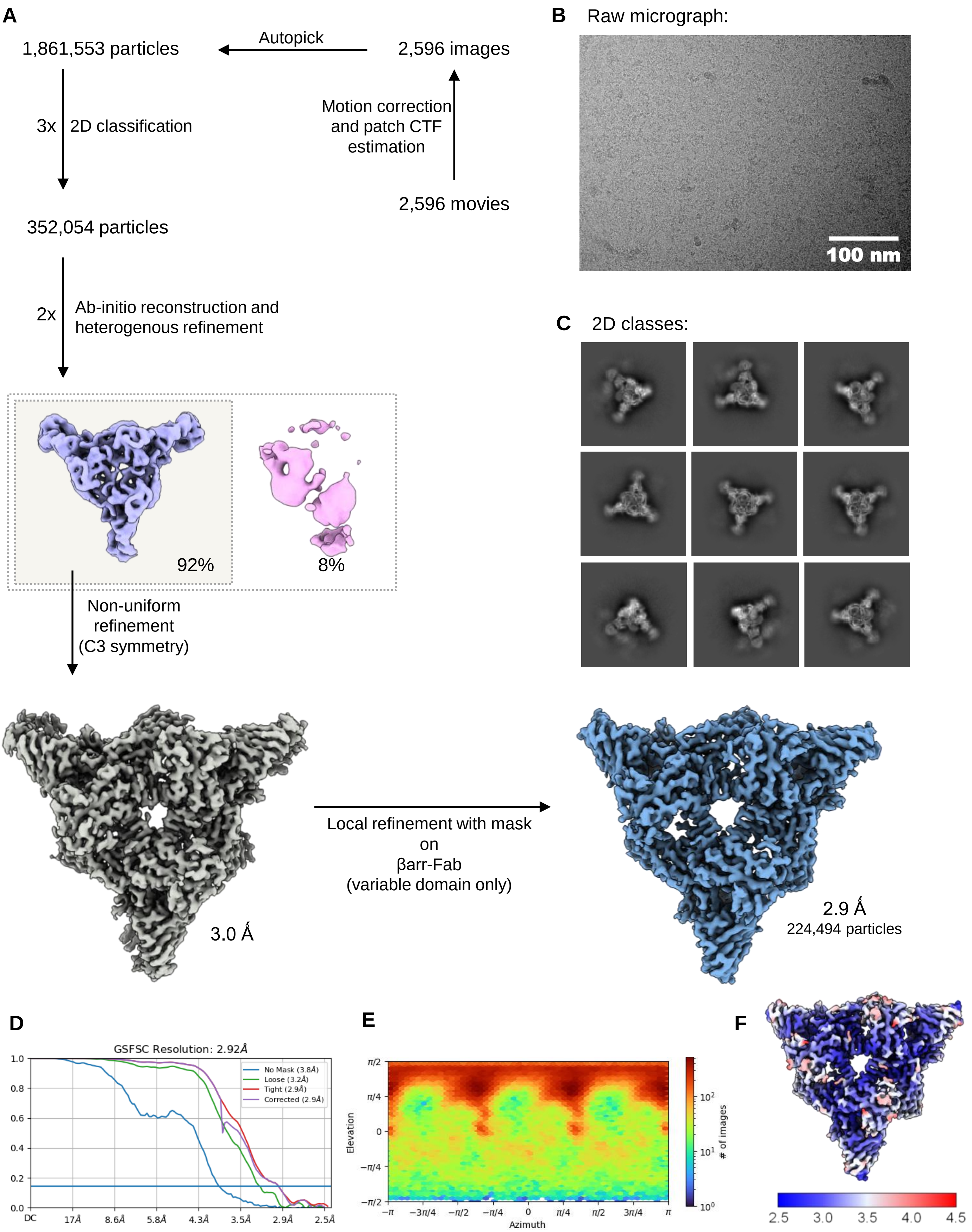
A
1,861,553 particles
Autopick
2,596 images
3x
2D classification
Motion correction and patch CTF estimation
2,596 movies
352,054 particles
2x
Ab-initio reconstruction and heterogenous refinement
92%
8%
Non-uniform refinement (C3 symmetry)
3.0 Å
Local refinement with mask on βarr-Fab (variable domain only)
2.9 Å
224,494 particles
B
Raw micrograph:
100 nm
C
2D classes:
D
GSFSC Resolution: 2.92Å
No Mask (3.8Å)
Loose (3.2Å)
Tight (2.9Å)
Corrected (2.9Å)
E
Elevation
Azimuth
### of images
F
2.5
3.0
3.5
4.0
4.5

**Figure S5. Cryo-EM reconstruction of the M2Rpp-βarr2-Fab30 complex.**

**(A)** Schematic representation of cryo-EM data processing pipeline. **(B)** Representative motion corrected micrograph. **(C)** Selected 2D class averages. **(D)** Gold standard fourier shell correlation curve (GFSC) at a threshold of 0.143 was used to determine the overall resolution of the map. **(E)** Angular plot of the particles used for 3D reconstruction. **(F)** Local resolution map of the 3D reconstruction in front view. (Scale bar: 100nm).

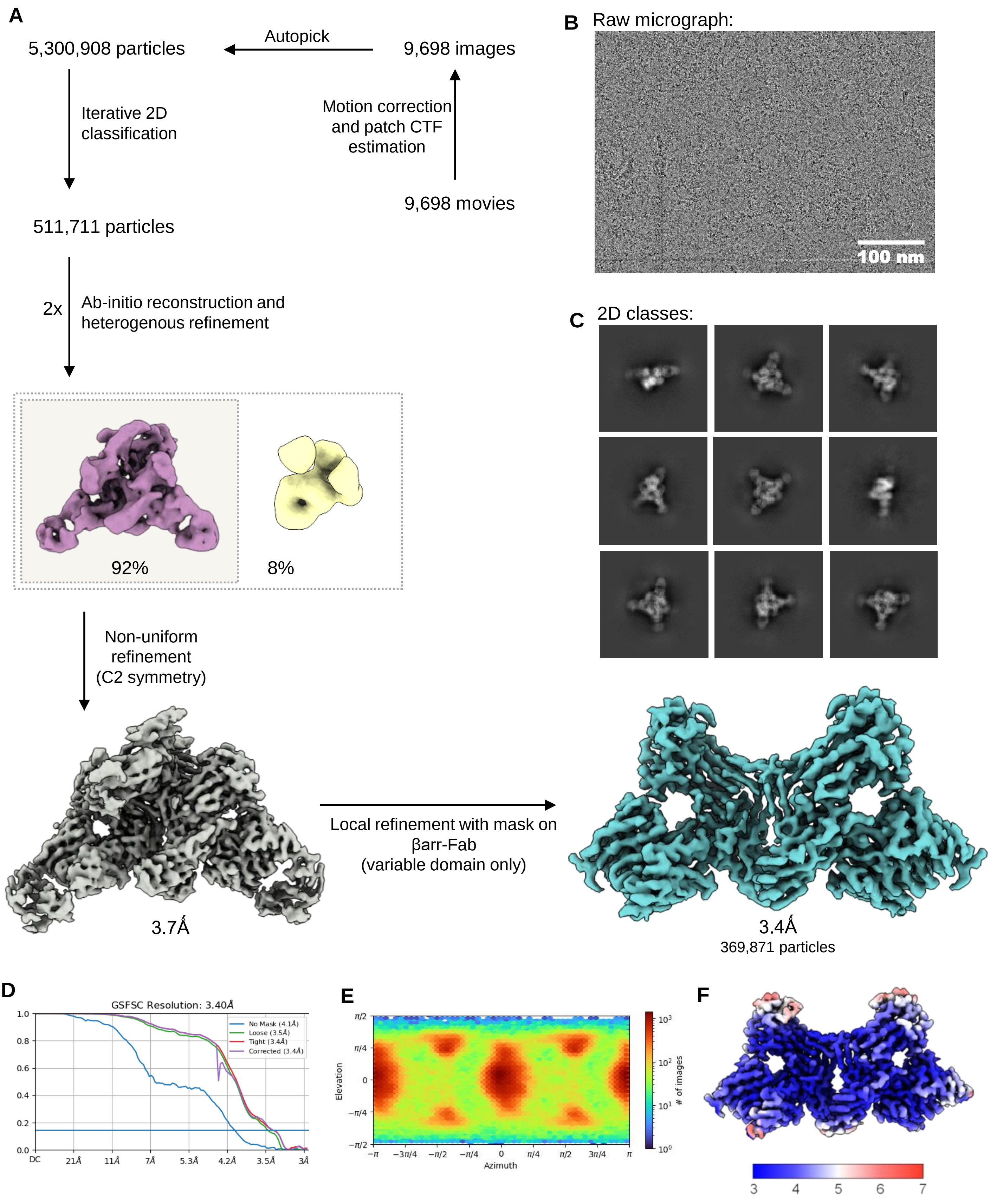

A
5,300,908 particles
Autopick
9,698 images
Iterative 2D classification
Motion correction and patch CTF estimation
9,698 movies
511,711 particles
2x
Ab-initio reconstruction and heterogenous refinement
92%
8%
Non-uniform refinement (C2 symmetry)
3.7Å
Local refinement with mask on βarr-Fab (variable domain only)
3.4Å
369,871 particles
B
Raw micrograph:
100 nm
C
2D classes:
D
GSFSC Resolution: 3.40Å
No Mask (4.1Å)
Loose (3.5Å)
Tight (3.4Å)
Corrected (3.4Å)
DC
21Å
11Å
7Å
5.3Å
4.2Å
3.5Å
3Å
E
Elevation
Azimuth
### of images
F
3
4
5
6
7

**Figure S6. Cryo-EM reconstruction of the D6Rpp-βarr1-Fab30 complex.**

**(A)** Schematic representation of cryo-EM data processing pipeline. **(B)** Representative motion corrected micrograph. **(C)** Selected 2D class averages. **(D)** Gold standard fourier shell correlation curve (GFSC) at 0.143 cut-off was used to determine the overall resolution of the map. **(E)** Angular distribution of the particles used for 3D reconstruction. **(F)** Local resolution map of the 3D reconstruction in front view. (Scale bar: 100nm).

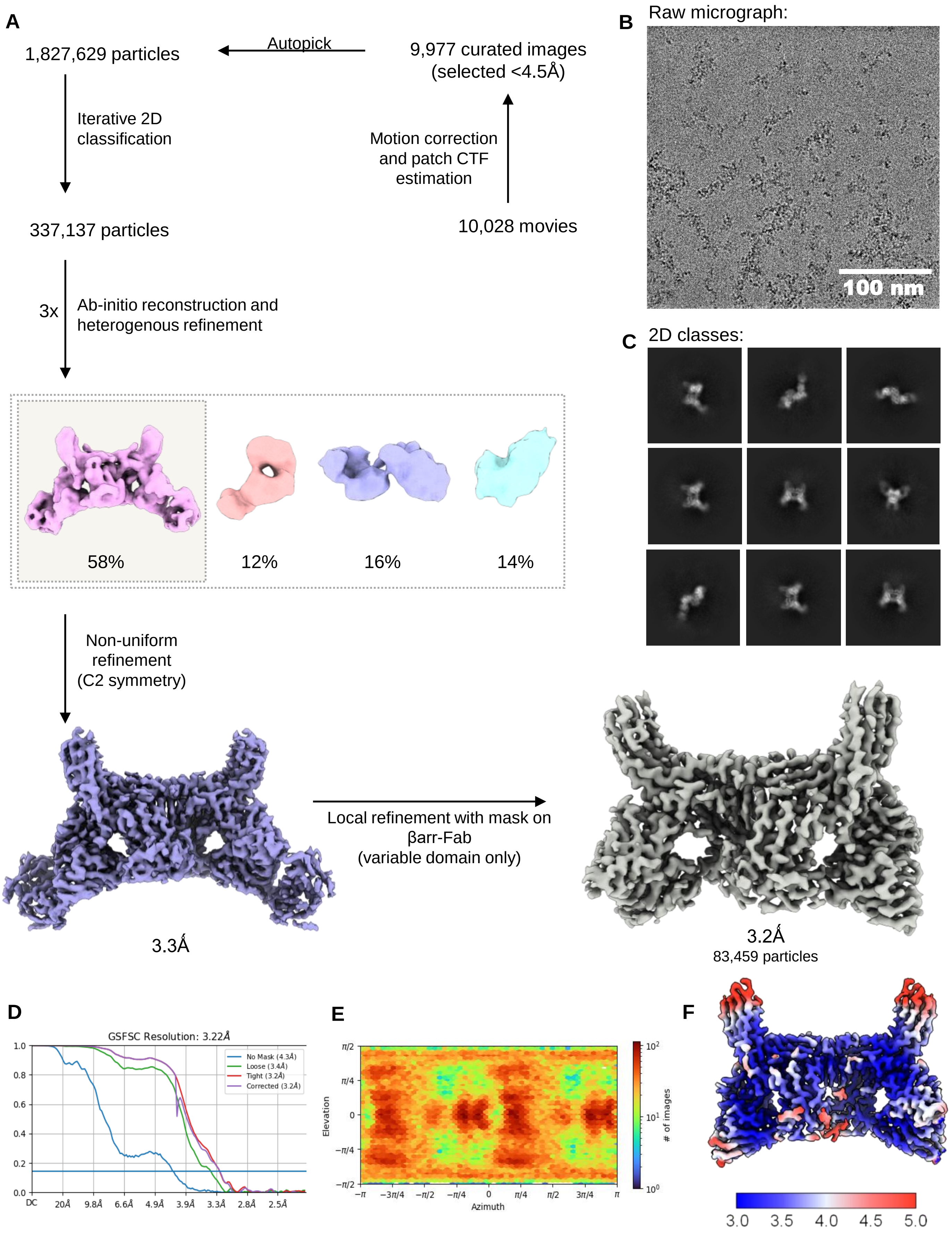
A
1,827,629 particles
Autopick
9,977 curated images
(selected <4.5Å)
Iterative 2D
classification
Motion correction
and patch CTF
estimation
337,137 particles
10,028 movies
3x
Ab-initio reconstruction and
heterogenous refinement
58%
12%
16%
14%
Non-uniform
refinement
(C2 symmetry)
3.3Å
Local refinement with mask on
βarr-Fab
(variable domain only)
3.2Å
83,459 particles
B
Raw micrograph:
100 nm
C
2D classes:
D
GSFSC Resolution: 3.22Å
No Mask (4.3Å)
Loose (3.4Å)
Tight (3.2Å)
Corrected (3.2Å)
DC
20Å
9.8Å
6.6Å
4.9Å
3.9Å
3.3Å
2.8Å
2.5Å
E
Elevation
Azimuth
### of images
F
3.0
3.5
4.0
4.5
5.0

**Figure S7. Cryo-EM reconstruction of the D6Rpp-βarr2-Fab30 complex.**

**(A)** Schematic representation of cryo-EM data processing pipeline. **(B)** Representative motion corrected micrograph. **(C)** Selected 2D class averages. **(D)** Gold standard fourier shell correlation curve (GFSC) at 0.143 cut-off was used to determine the overall resolution of the map. **(E)** Angular plot showing distribution of the particles used for 3D reconstruction. **(F)** Local resolution map of the 3D reconstruction in front view. (Scale bar: 100nm).

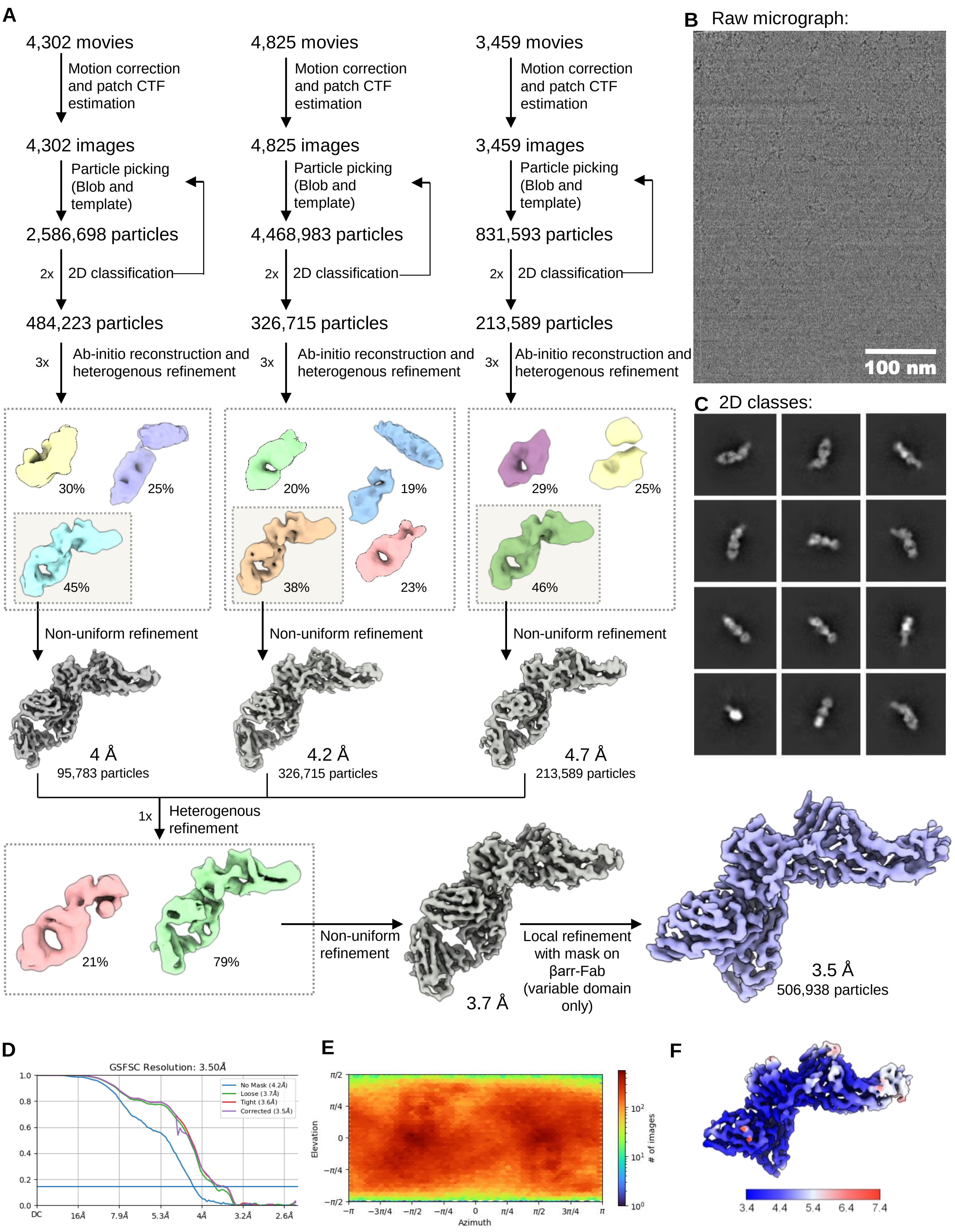

A
4,302 movies
Motion correction and patch CTF estimation
4,302 images
Particle picking (Blob and template)
2,586,698 particles
2x 2D classification
484,223 particles
3x Ab-initio reconstruction and heterogenous refinement
30%
25%
45%
Non-uniform refinement
4 Å
95,783 particles
4,825 movies
Motion correction and patch CTF estimation
4,825 images
Particle picking (Blob and template)
4,468,983 particles
2x 2D classification
326,715 particles
3x Ab-initio reconstruction and heterogenous refinement
20%
19%
38%
23%
Non-uniform refinement
4.2 Å
326,715 particles
3,459 movies
Motion correction and patch CTF estimation
3,459 images
Particle picking (Blob and template)
831,593 particles
2x 2D classification
213,589 particles
3x Ab-initio reconstruction and heterogenous refinement
29%
25%
46%
Non-uniform refinement
4.7 Å
213,589 particles
1x Heterogenous refinement
21%
79%
Non-uniform refinement
3.7 Å
Local refinement with mask on βarr-Fab (variable domain only)
3.5 Å
506,938 particles
B Raw micrograph:
100 nm
C 2D classes:
D
GSFSC Resolution: 3.50Å
No Mask (4.2Å)
Loose (3.7Å)
Tight (3.6Å)
Corrected (3.5Å)
DC
16Å
7.9Å
5.3Å
4Å
3.2Å
2.6Å
E
Elevation
Azimuth
### of images
F
3.4
4.4
5.4
6.4
7.4

**Figure S8. Cryo-EM reconstruction of the basal βarr2.**

**(A)** Schematic representation of the cryo-EM data processing pipeline. **(B)** Representative motion corrected micrograph. **(C)** Selected 2D class averages. **(D)** Gold standard fourier shell correlation curve (GFSC) at 0.143 cut-off was used to determine the overall resolution of the map. **(E)** Angular plot of the particles used for 3D reconstruction. **(F)** Local resolution map of the 3D reconstruction in front view. (Scale bar: 100nm).

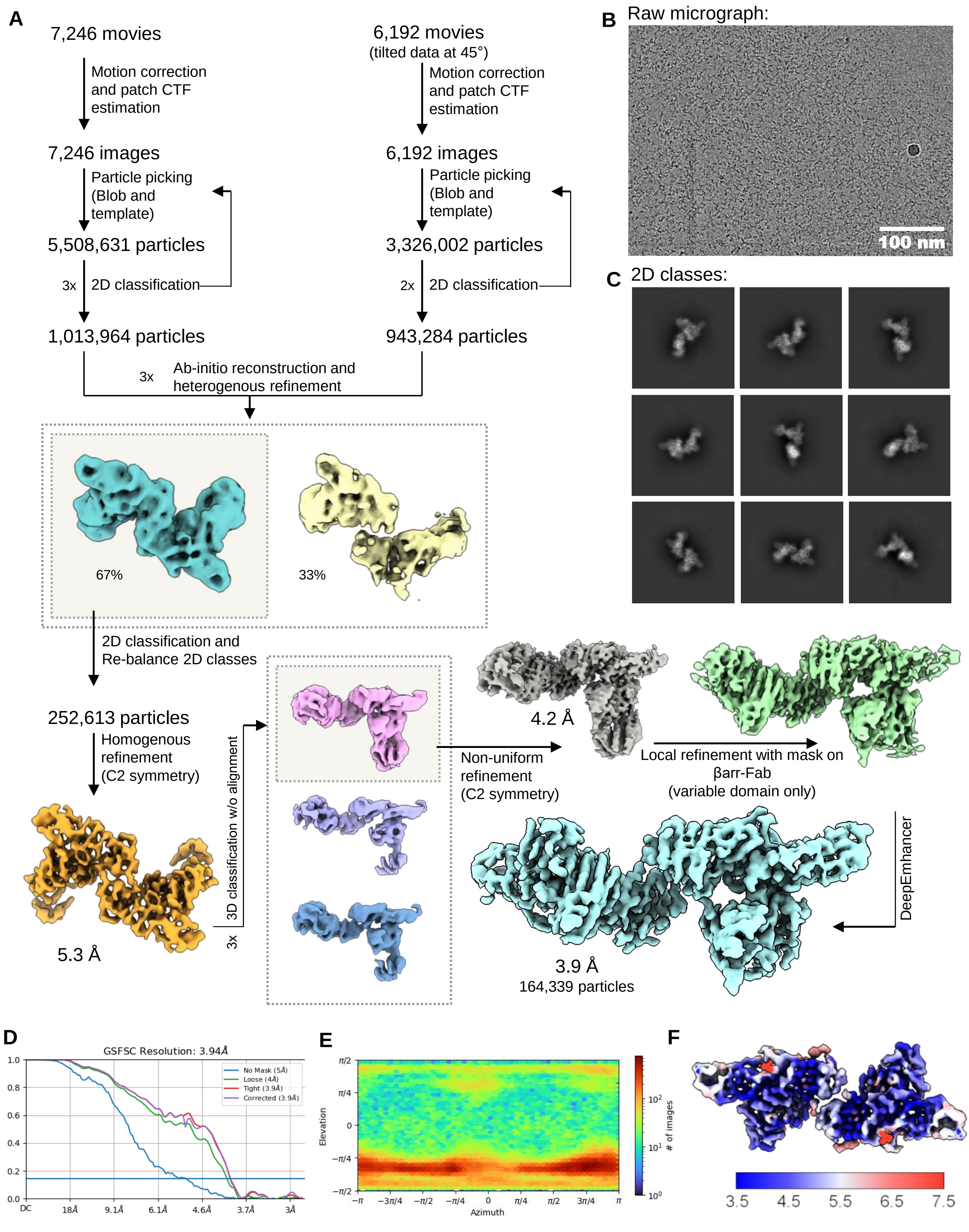
A
7,246 movies
Motion correction and patch CTF estimation
7,246 images
Particle picking (Blob and template)
5,508,631 particles
3x
2D classification
1,013,964 particles
6,192 movies
(tilted data at 45°)
Motion correction and patch CTF estimation
6,192 images
Particle picking (Blob and template)
3,326,002 particles
2x
2D classification
943,284 particles
3x
Ab-initio reconstruction and heterogenous refinement
67%
33%
2D classification and Re-balance 2D classes
252,613 particles
Homogenous refinement (C2 symmetry)
5.3 Å
3D classification w/o alignment
3x
Non-uniform refinement (C2 symmetry)
4.2 Å
Local refinement with mask on βarr-Fab (variable domain only)
DeepEmhancer
3.9 Å
164,339 particles
B
Raw micrograph:
100 nm
C
2D classes:
D
GSFSC Resolution: 3.94Å
No Mask (5Å)
Loose (4Å)
Tight (3.9Å)
Corrected (3.9Å)
E
Elevation
Azimuth
### of images
F
3.5
4.5
5.5
6.5
7.5

**Figure S9. Cryo-EM reconstruction of the C3aRpp-βarr1-Fab30 complex.**

**(A)** Schematic representation of cryo-EM data processing pipeline. **(B)** Representative motion corrected micrograph. **(C)** Selected 2D class averages. **(D)** Gold standard fourier shell correlation curve (GFSC) at 0.143 cut-off was used to determine the overall resolution of the map. **(E)** Angular plot of the particles used for 3D reconstruction. **(F)** Local resolution map of the 3D reconstruction in front view. (Scale bar: 100nm).

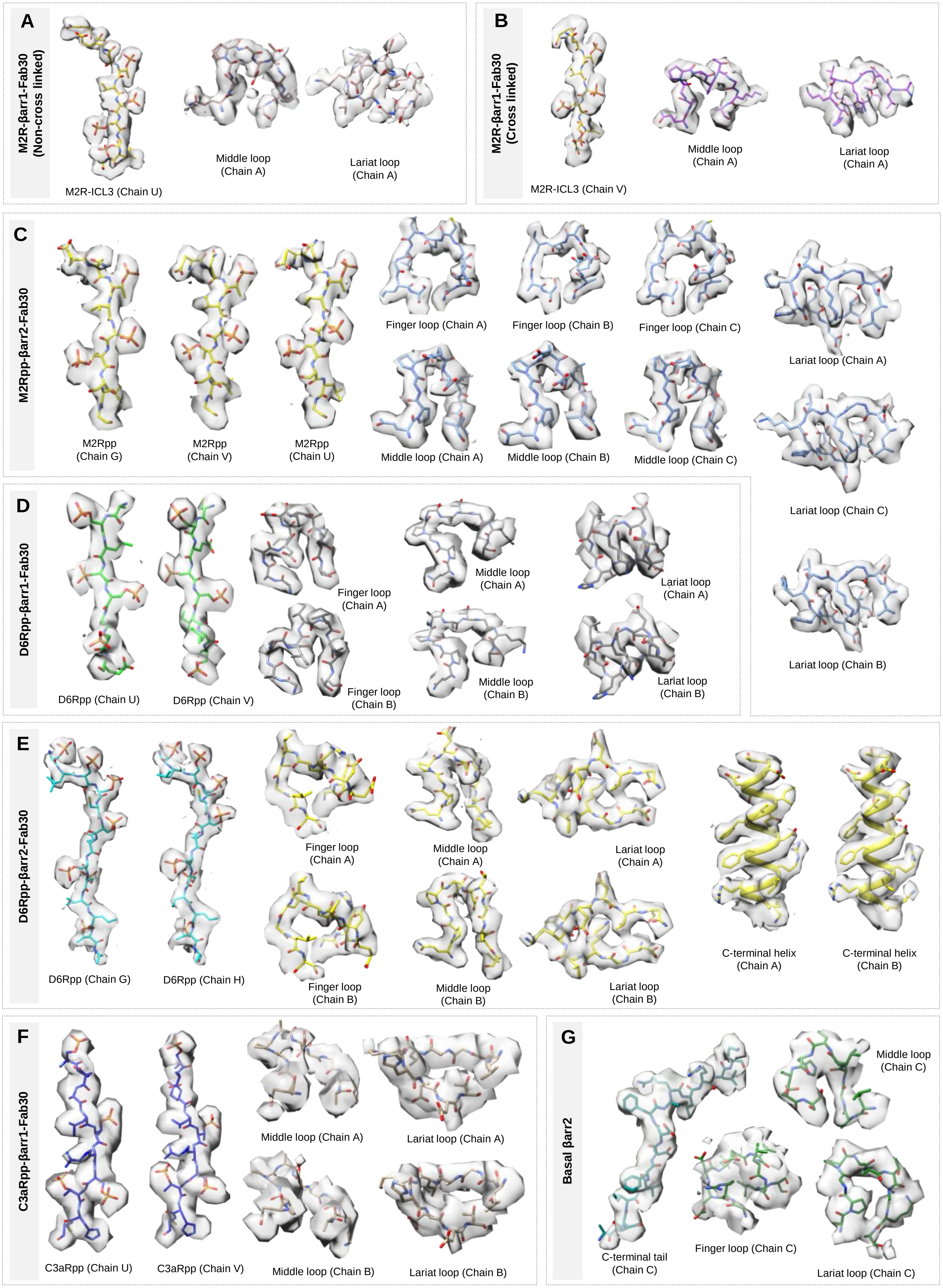

A
M2R-βarr1-Fab30 (Non-cross linked)
M2R-ICL3 (Chain U)
Middle loop (Chain A)
Lariat loop (Chain A)
B
M2R-βarr1-Fab30 (Cross linked)
M2R-ICL3 (Chain V)
Middle loop (Chain A)
Lariat loop (Chain A)
C
M2Rpp-βarr2-Fab30
M2Rpp (Chain G)
M2Rpp (Chain V)
M2Rpp (Chain U)
Finger loop (Chain A)
Finger loop (Chain B)
Finger loop (Chain C)
Middle loop (Chain A)
Middle loop (Chain B)
Middle loop (Chain C)
Lariat loop (Chain A)
Lariat loop (Chain C)
Lariat loop (Chain B)
D
D6Rpp-βarr1-Fab30
D6Rpp (Chain U)
D6Rpp (Chain V)
Finger loop (Chain A)
Finger loop (Chain B)
Middle loop (Chain A)
Middle loop (Chain B)
Lariat loop (Chain A)
Lariat loop (Chain B)
E
D6Rpp-βarr2-Fab30
D6Rpp (Chain G)
D6Rpp (Chain H)
Finger loop (Chain A)
Finger loop (Chain B)
Middle loop (Chain A)
Middle loop (Chain B)
Lariat loop (Chain A)
Lariat loop (Chain B)
C-terminal helix (Chain A)
C-terminal helix (Chain B)
F
C3aRpp-βarr1-Fab30
C3aRpp (Chain U)
C3aRpp (Chain V)
Middle loop (Chain A)
Middle loop (Chain B)
Lariat loop (Chain A)
Lariat loop (Chain B)
G
Basal βarr2
C-terminal tail (Chain C)
Finger loop (Chain C)
Middle loop (Chain C)
Lariat loop (Chain C)

**Figure S10. Exemplary electron density maps.**

EM densities for the phosphopeptides, finger loops, middle loops, lariat loops and helices in the **(A)** M2R-βarr1-Fab30 (non-cross linked) complex, **(B)** M2R-βarr1-Fab30 (cross linked) complex, **(C)** M2Rpp-βarr2-Fab30 complex, **(D)** D6Rpp-βarr1-Fab30 complex, **(E)** D6Rpp-βarr2-Fab30 complex, **(F)** C3aRpp-βarr1-Fab30 complex, and **(G)** Basal βarr2 structure.

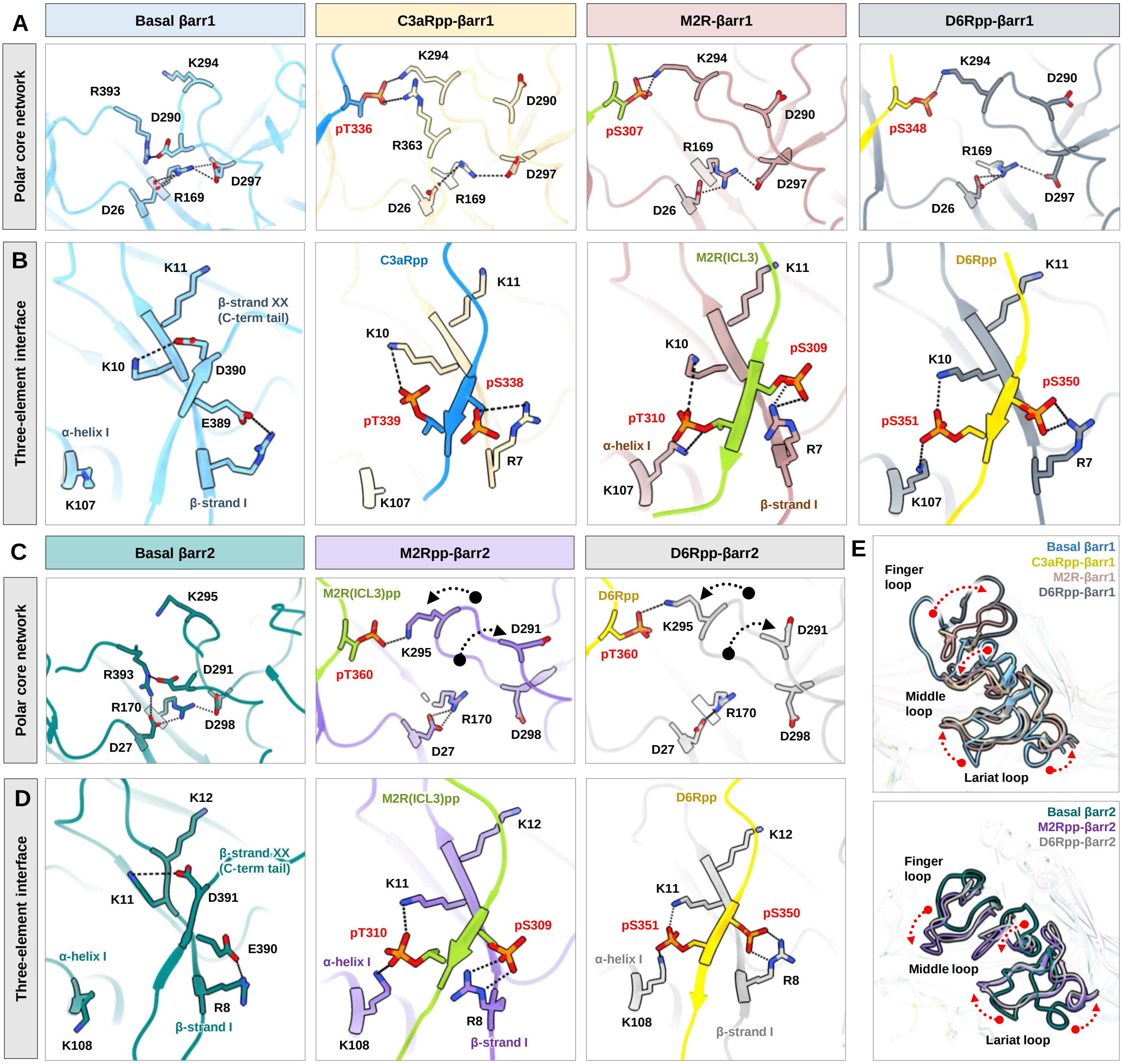
A
Basal βarr1
C3aRpp-βarr1
M2R-βarr1
D6Rpp-βarr1
Polar core network
K294
R393
D290
D297
R169
D26
pT336
R363
pS307
pS348
B
Three-element interface
K11
K10
β-strand XX (C-term tail)
D390
E389
α-helix I
K107
β-strand I
C3aRpp
pS338
pT339
R7
M2R(ICL3)
pS309
pT310
D6Rpp
pS350
pS351
C
Basal βarr2
M2Rpp-βarr2
D6Rpp-βarr2
K295
D291
R170
D298
D27
M2R(ICL3)pp
pT360
D
K12
D391
E390
R8
K108
E
Basal βarr1
C3aRpp-βarr1
M2R-βarr1
D6Rpp-βarr1
Finger loop
Middle loop
Lariat loop
Basal βarr2
M2Rpp-βarr2
D6Rpp-βarr2

**Figure S11. Structural signature hallmarks of active βarr1/2.**

**(A)** Polar-core network disruption in all solved structures of βarr1 are shown (basal state βarr1, PDB 1G4M). **(B)** Key residues involved in the three-element interface of βarr1 have been highlighted. **(C)** Polar-core network disruption in all solved structures of βarr2 are shown. **(D)** Key residues involved in the three-element interface have been highlighted (the structure of basal state βarr2 used here is the one that has been determined in this study). **(E)** The phosphorylated residues in GPCRs adopt a "PXPP" type motif for activation of βarrs. Structural alignment has been carried out on the highest resolution structure, M2Rpp-βarr2.

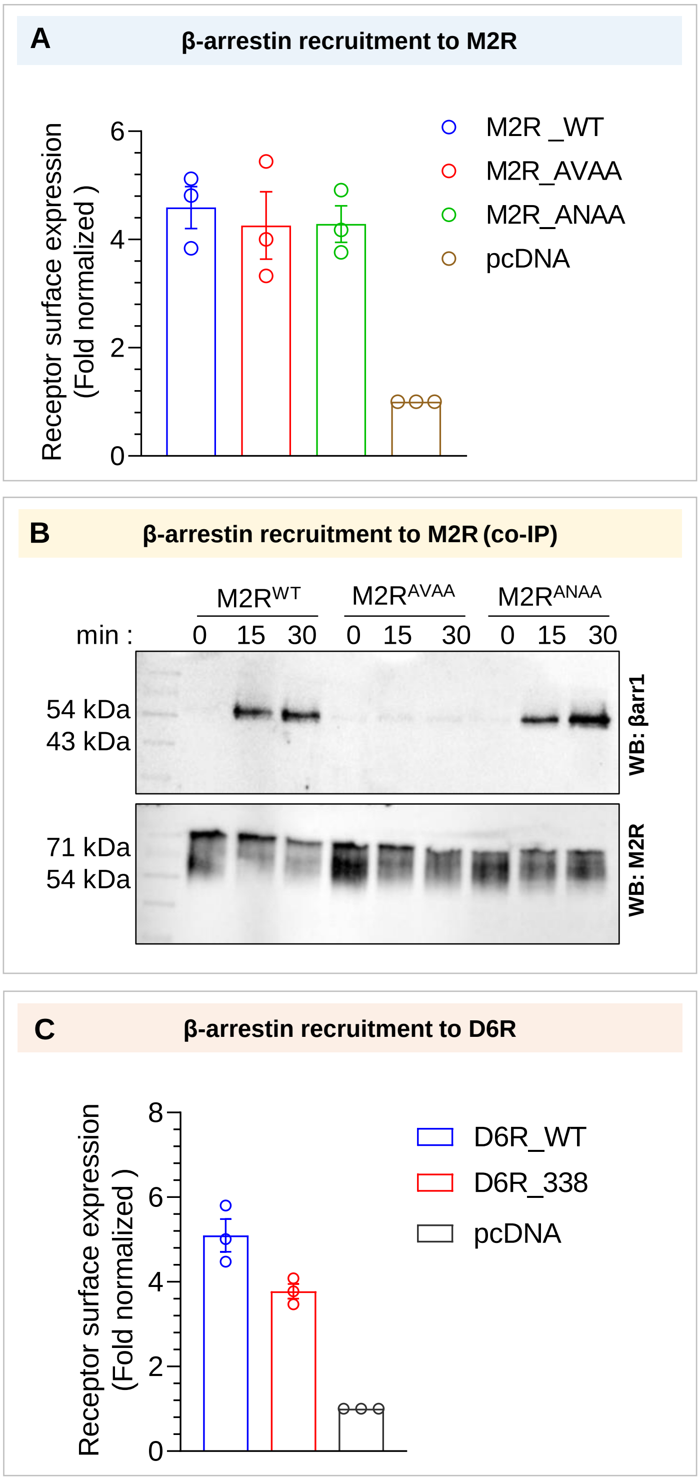
A
β-arrestin recruitment to M2R
Receptor surface expression
(Fold normalized )
0
2
4
6
M2R _WT
M2R_AVAA
M2R_ANAA
pcDNA
B
β-arrestin recruitment to M2R (co-IP)
M2RWT
M2RAVAA
M2RANAA
min :
0 15 30 0 15 30 0 15 30
54 kDa
43 kDa
WB: βarr1
71 kDa
54 kDa
WB: M2R
C
β-arrestin recruitment to D6R
Receptor surface expression
(Fold normalized )
0
2
4
6
8
D6R_WT
D6R_338
pcDNA

**Figure S12. Surface expression profiles of receptors in βarr1 recruitment assays.**

**(A)** Surface expression of $M2R^{WT}$, and $M2R^{Mutants}$ in the NanoBiT assay was measured by using whole cell ELISA. **(B)** Role of mutants in βarr recruitment is further assessed by co-immunoprecipitation assay. On Carbachol stimulation, $M2R^{AVAA}$ showed dramatic reduction in βarr1 recruitment. A representative blot and densitometry-based quantification (mean±SEM; n=4; normalized with M2R 30 min stimulation condition signal as 100%; Two-way ANOVA, Tukey's multiple comparisons test) is presented (p values). **(C)** Surface expression of $D6R^{WT}$, and $D6R^{Mutants}$ in the NanoBiT assays was measured by using whole cell ELISA.

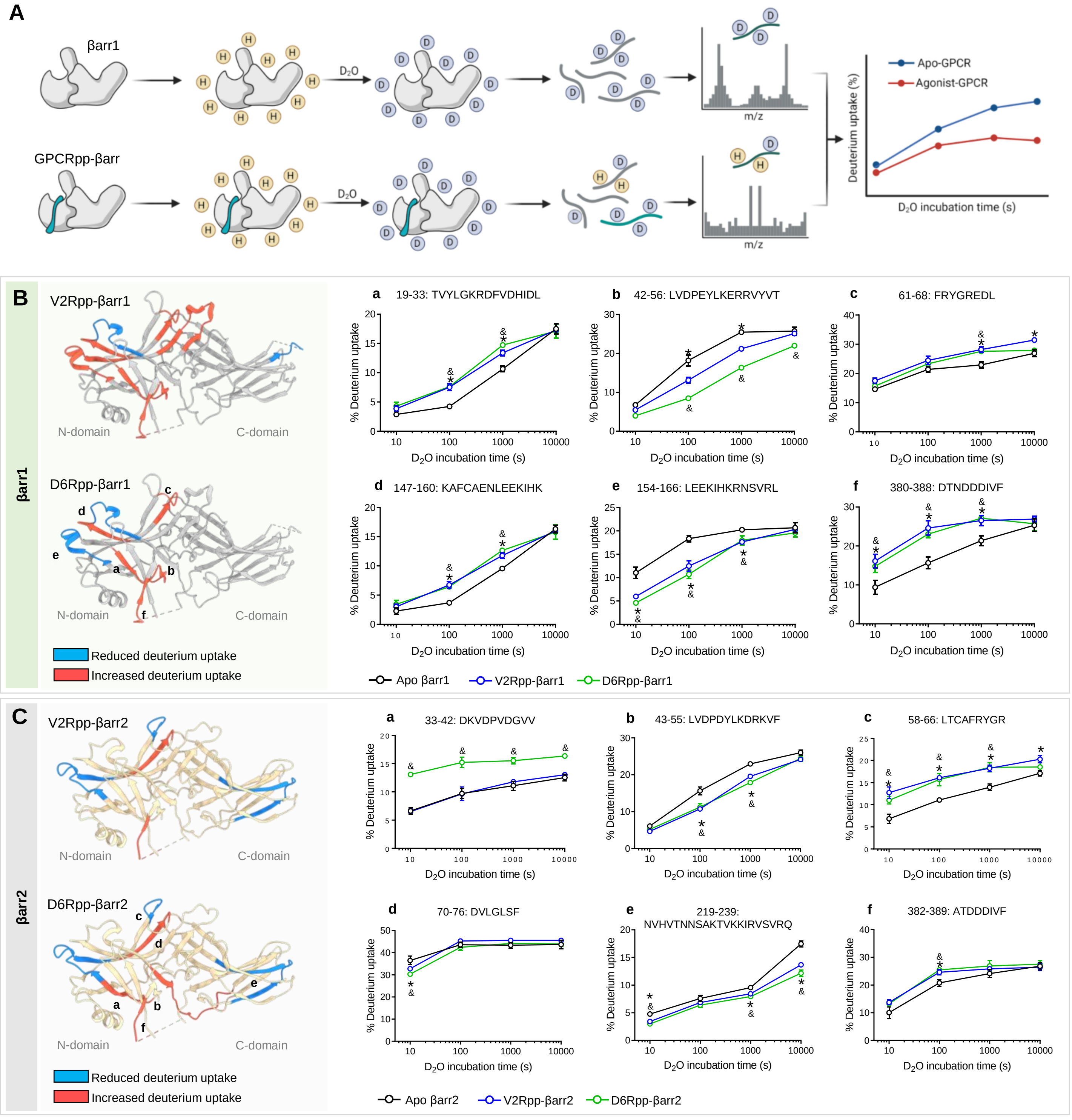
A
βarr1
GPCRpp-βarr
D2O
m/z
Apo-GPCR
Agonist-GPCR
Deuterium uptake (%)
D2O incubation time (s)
B
βarr1
V2Rpp-βarr1
D6Rpp-βarr1
N-domain
C-domain
Reduced deuterium uptake
Increased deuterium uptake
a 19-33: TVYLGKRDFVDHIDL
b 42-56: LVDPEYLKERRVYVT
c 61-68: FRYGREDL
d 147-160: KAFCAENLEEKIHK
e 154-166: LEEKIHKRNSVRL
f 380-388: DTNDDDIVF
% Deuterium uptake
D2O incubation time (s)
Apo βarr1
V2Rpp-βarr1
D6Rpp-βarr1
C
βarr2
V2Rpp-βarr2
D6Rpp-βarr2
a 33-42: DKVDPVDGVV
b 43-55: LVDPDYLKDRKVF
c 58-66: LTCAFRYGR
d 70-76: DVLGLSF
e 219-239: NVHVTNNSAKTVKKIRVSVRQ
f 382-389: ATDDDIVF
Apo βarr2
V2Rpp-βarr2
D6Rpp-βarr2

**Figure S13. HDX-MS analysis of βarr activation by phosphopeptides.**

**(A**) Schematic representation of working principle of HDX-MS. **(B, C)** Deuterium exchange plots of V2Rpp and D6Rpp activated βarr1 and βarr2, respectively.

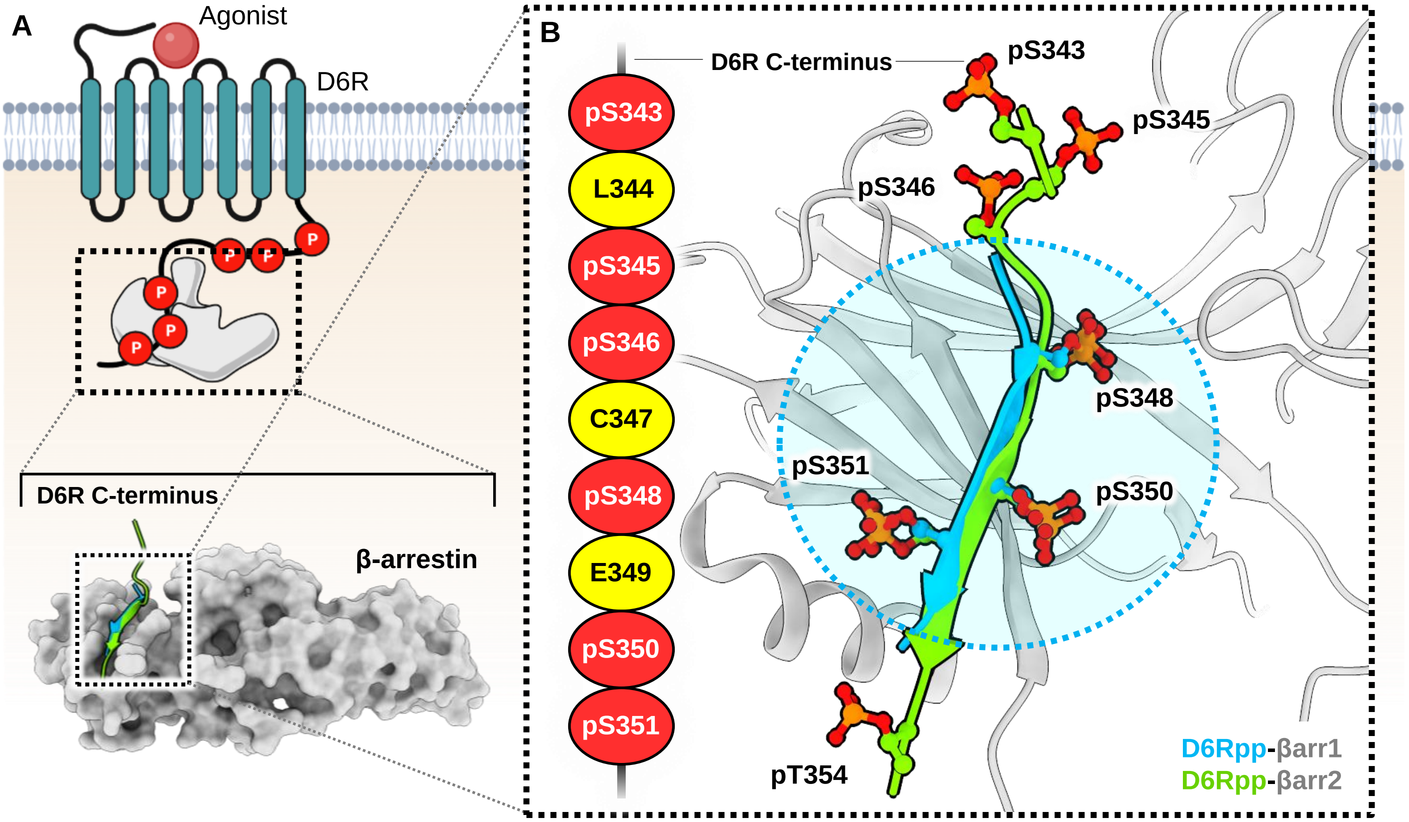
A
Agonist
D6R
P
D6R C-terminus
β-arrestin
B
D6R C-terminus
pS343
L344
pS345
pS346
C347
pS348
E349
pS350
pS351
pS343
pS345
pS346
pS348
pS351
pS350
pT354
D6Rpp-βarr1
D6Rpp-βarr2

**Figure S14. Selectivity of phosphorylation residues for βarr binding on the C-terminus of D6R.**

**(A)** A cartoon representation of D6R showing βarr engagement to the phosphorylated C-terminus (top). Structural snapshot of D6Rpp-βarr2 in surface representation highlighting the position of D6Rpp in both βarr1 (cyan) and βarr2 (green). **(B)** The stretch of amino acids in the C-terminus of D6R harboring phosphorylation clusters have been depicted (left). Among the clusters $^{343}$pS-L-pS-pS$^{346}$ and $^{348}$pS-E-pS-pS$^{351}$, the $^{348}$pS-E-pS-pS$^{351}$ cluster (dotted cyan circle) is found to be preferred for interaction and subsequent activation with both βarr1 and βarr2.

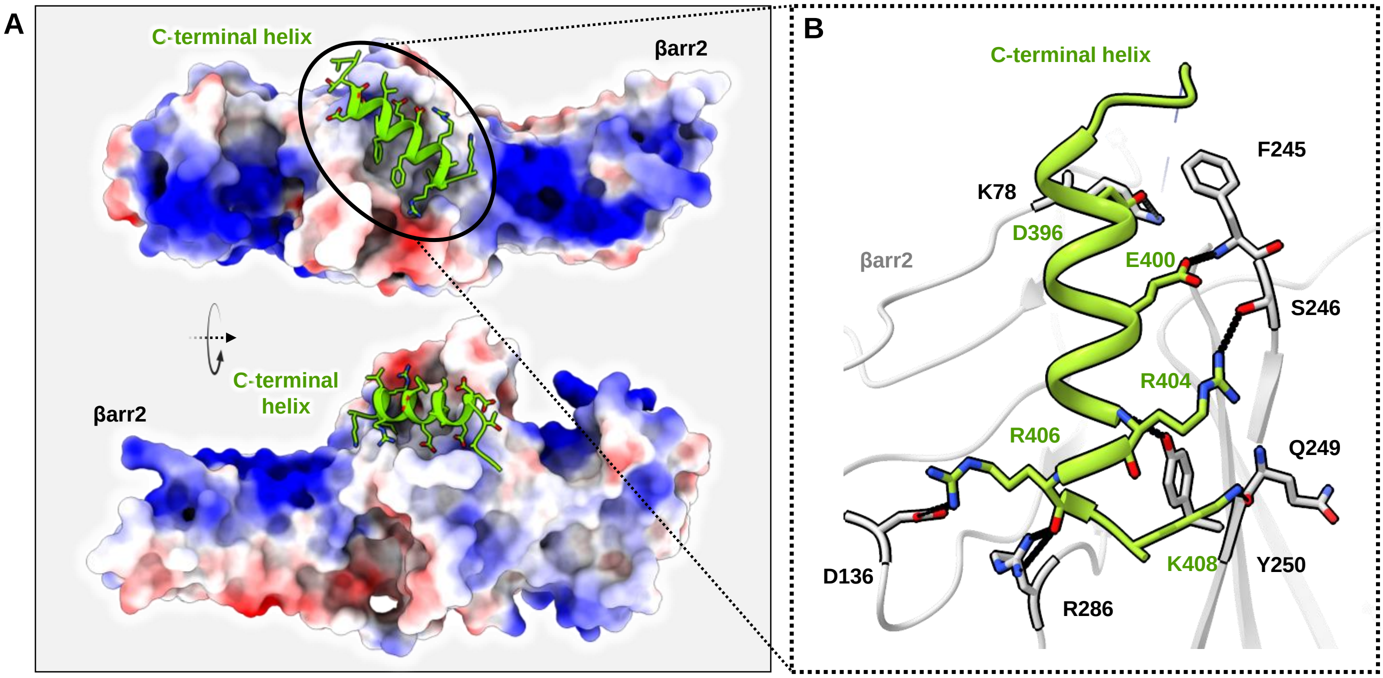
A
C-terminal helix
βarr2
C-terminal
helix
βarr2
B
C-terminal helix
F245
K78
D396
βarr2
E400
S246
R404
R406
Q249
D136
K408
Y250
R286

**Figure S15. Binding interface of C-terminal helix on βarr2.**

**(A)** Binding groove of the C-terminal helix on βarr2 (central crest) shown as charged surface. **(B)** Residue-residue contacts between the C-terminal helix and βarr2 are presented.

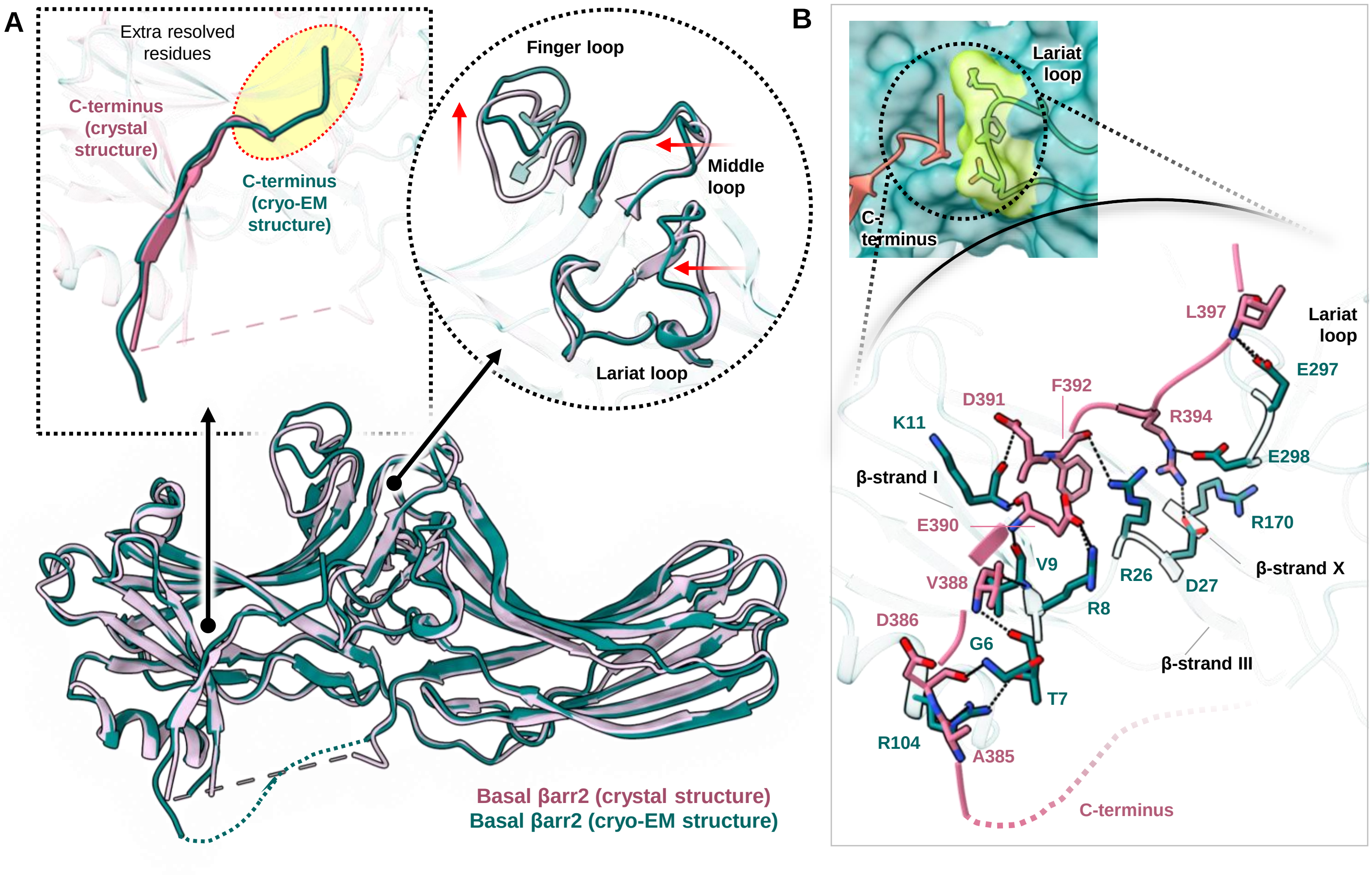

A
Extra resolved residues
C-terminus (crystal structure)
C-terminus (cryo-EM structure)
Finger loop
Middle loop
Lariat loop
Basal βarr2 (crystal structure)
Basal βarr2 (cryo-EM structure)
B
Lariat loop
C-terminus
L397
Lariat loop
E297
F392
D391
R394
K11
E298
β-strand I
R170
E390
β-strand X
V9
R26
D27
V388
R8
D386
G6
β-strand III
T7
R104
A385
C-terminus

**Figure S16. Structural comparison of basal βarr2 determined through x-ray crystallography and cryo-EM.**

**(A)** Structural superimposition of basal βarr2 crystal structure with basal βarr2 cryo-EM structure (middle). Only the C-terminus has been highlighted to show the extra residues resolved in the cryo-EM structure (left). Conformational changes in the loops of βarr2 have been illustrated by comparing both structures (right). **(B)** Interaction of the C-terminal residues of basal βarr2 with the lariat loop residues. βarr2 in surface and C-terminal tail in ribbon representation (top). Residue contacts between the C-terminus and various regions of βarr2 (bottom).

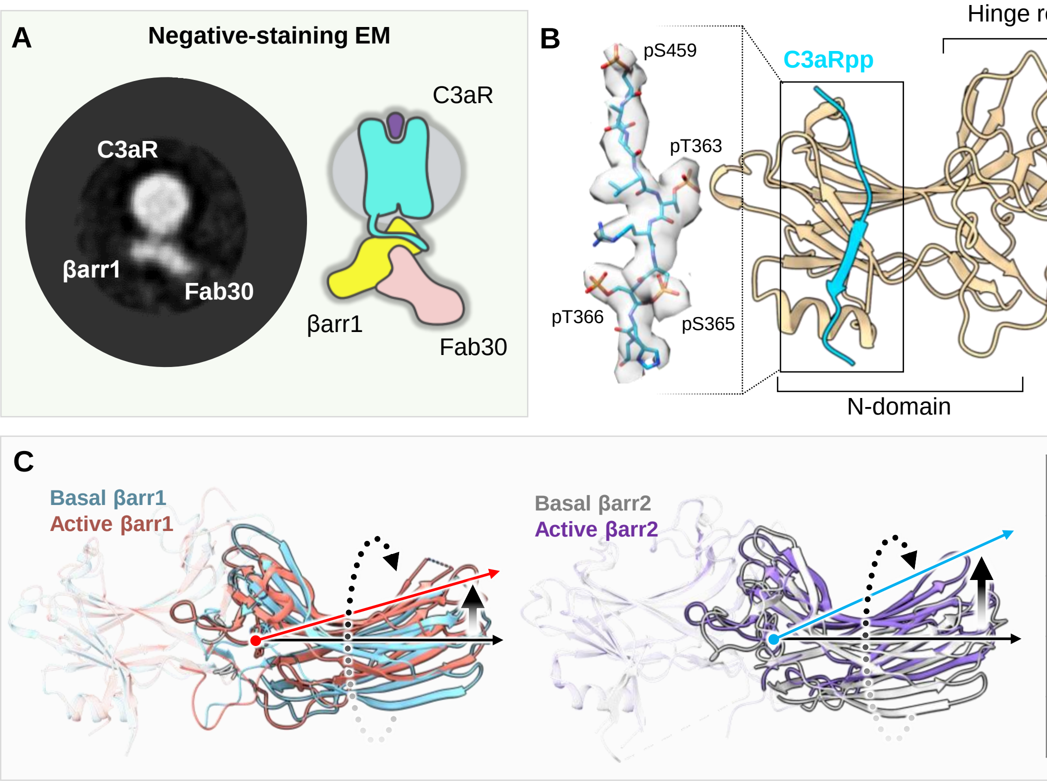


| Receptor | βarr1 | βarr2 |
|---|---|---|
| V2R | ~21° | ~25° |
| C5aR1 | ~20° | ~24° |
| M2R | ~18° | ~23° |
| D6R | ~20° | ~22° |

**Figure S17. Structure of C3aRpp-βarr1 complex and the differences in the strength of activation between two βarr isoforms.**

(**A**) Reconstitution of C3aR-βarr1 complex and visualization through negative staining EM. A cartoon representation to highlight the "hanging" mode of βarr1 complex organization. (**B**) Structure of C3aR phosphopeptide bound βarr1 complex in ribbon representation. EM density of C3aRpp is provided to the left. βarr1 attains active conformation with a C-domain rotation of 18.9°. **(C)** Structural superimposition of βarr1 and βarr2 in basal states and respective active states. Basal state βarr1 (PDB 1G4M), active state βarr1 (M2R-βarr1, non-cross linked structure) (left), Basal state βarr2 (cryo-EM βarr2 structure) and active state βarr2 (M2Rpp-βarr2) (middle), A table showing comparison of inter-domain rotation of βarr pairs upon activation by receptors.

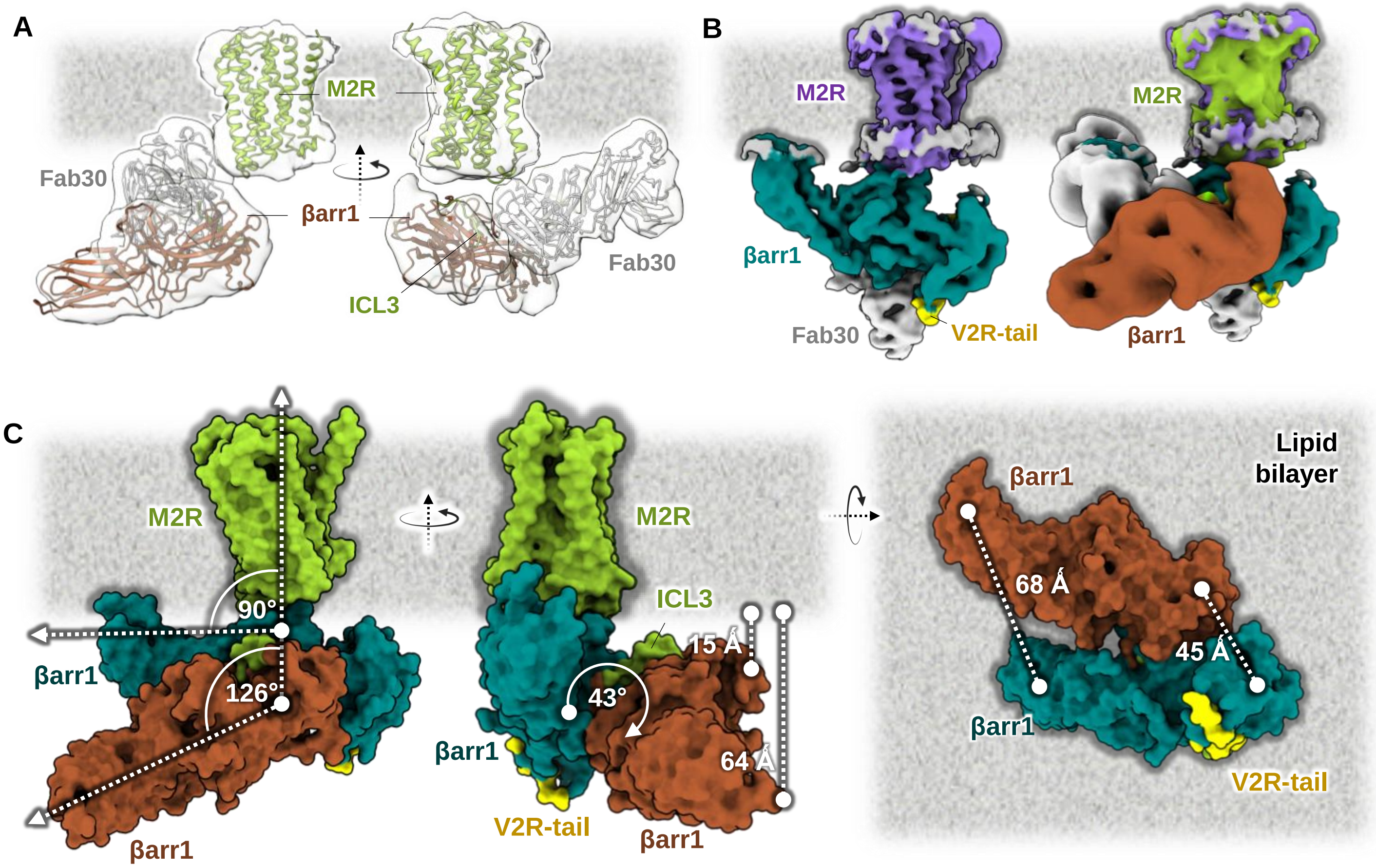
A
M2R
Fab30
βarr1
ICL3
Fab30
B
M2R
M2R
βarr1
Fab30
V2R-tail
βarr1
C
M2R
90°
βarr1
126°
βarr1
M2R
ICL3
15 Å
43°
βarr1
64 Å
V2R-tail
βarr1
Lipid
bilayer
βarr1
68 Å
45 Å
βarr1
V2R-tail

**Figure S18. Structural features of “hanging” M2R-βarr1 complex.**

**(A)** Cryo-EM 3D reconstruction of “hanging” M2R-βarr1-Fab30 complex. Density map representation (top) and map with fitted models (bottom). **(B)** Structural alignment of the receptor with previously determined cryo-EM map of M2V2R-βarr1 complex (PDB 6U1N) to illustrate the relative position of βarr1. **(C)** Aligning the receptor with the membrane axis reveals the position of βarr1 with respect to the lipid bilayer and the receptor. Structural superimposition was carried out on M2R (green; native M2R from PDB 6U1N, brown: βarr1 bound to ICL3 of M2R, teal: βarr1 from PDB 6U1N) to investigate the relative positions of βarr1.

A

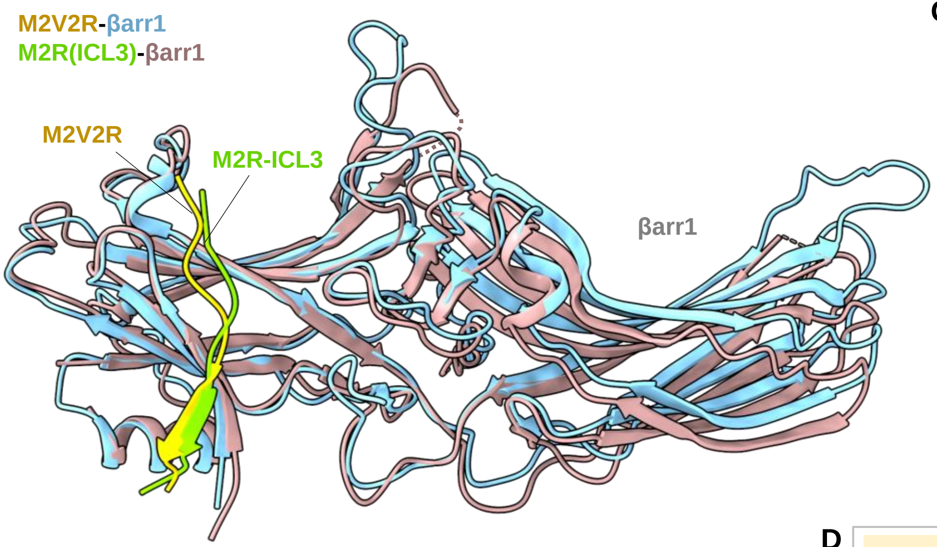


C

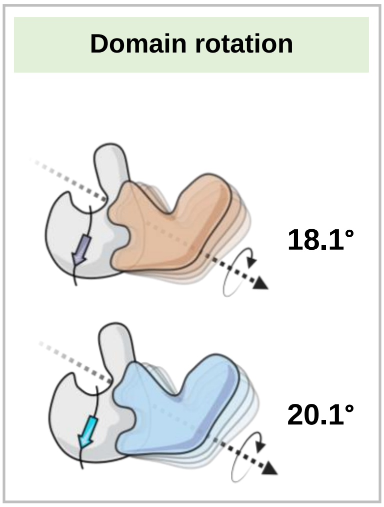


B **Phospho-residue contacts:**

| M2R-βarr1 | |
|---|---|
| Phospho-residue | βarr1 residue |
| pT307 | K11,R25,K294 |
| pS309 | R7 |
| pT310 | K10,K107 |

| M2V2R-βarr1 | |
|---|---|
| Phospho-residue | βarr1 residue |
| pS488 (pS357) | R165 |
| pT491 (pT359) | K11,R25,K294 |
| pS493 (pT361) | R7 |
| pS494 (pT362) | K10,K107 |

D

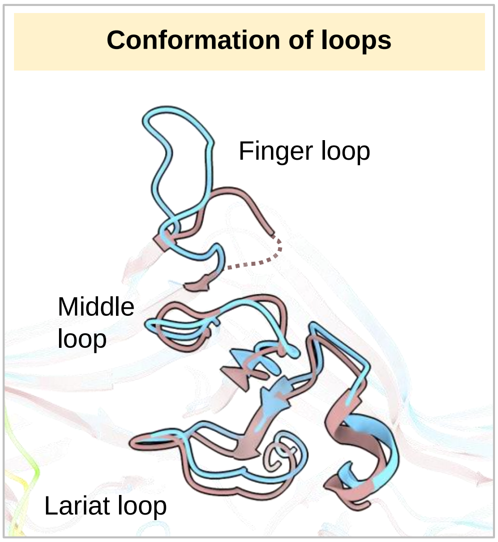

**Figure S19. Structural comparison of βarr conformation bound to M2V2R and M2R.**

**(A)** Superimposition of M2V2R-βarr1 (PDB 6U1N) and M2R-βarr1 on βarr1. Structures are shown in ribbon representation. **(B)** List of residue-residue contacts between the phosphorylated C-terminus or ICL3 in M2R-βarr1 (left) and M2V2R-βarr1 (right), respectively. The corresponding phospho-residue numbers in V2R have been mentioned below in every respective cell. **(C)** A comparison of C-domain rotation values compared to respective N-domains of M2R-βarr1 (top) and M2V2R-βarr1 (bottom). **(D)** Changes in conformations of key loops in βarr1 have been illustrated.

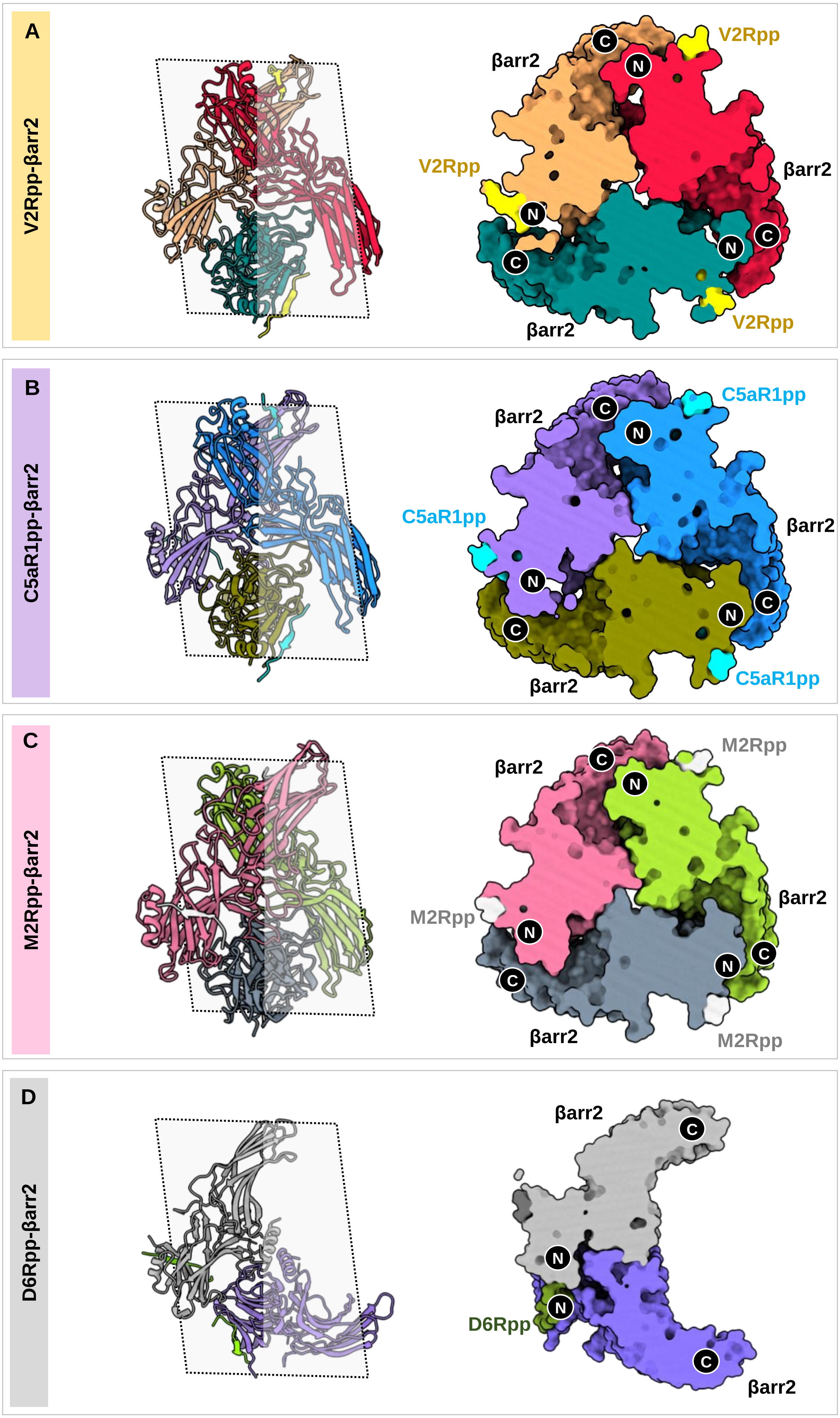
A
V2Rpp-βarr2
V2Rpp
βarr2
βarr2
V2Rpp
βarr2
V2Rpp
B
C5aR1pp-βarr2
C5aR1pp
βarr2
C5aR1pp
βarr2
βarr2
C5aR1pp
C
M2Rpp-βarr2
βarr2
M2Rpp
βarr2
M2Rpp
βarr2
M2Rpp
D
D6Rpp-βarr2
βarr2
D6Rpp
βarr2

**Figure S20. Oligomerization states of βarr2.**

**(A, B, C, D)** Oligomerization mode and domain organization of different βarr2 oligomers in V2Rpp-βarr2, C5aR1pp-βarr2, M2Rpp-βarr2 and D6R-βarr2 structures, respectively. Oligomers have been shown in surface slice representation (right) with the defined axis of slice (left).

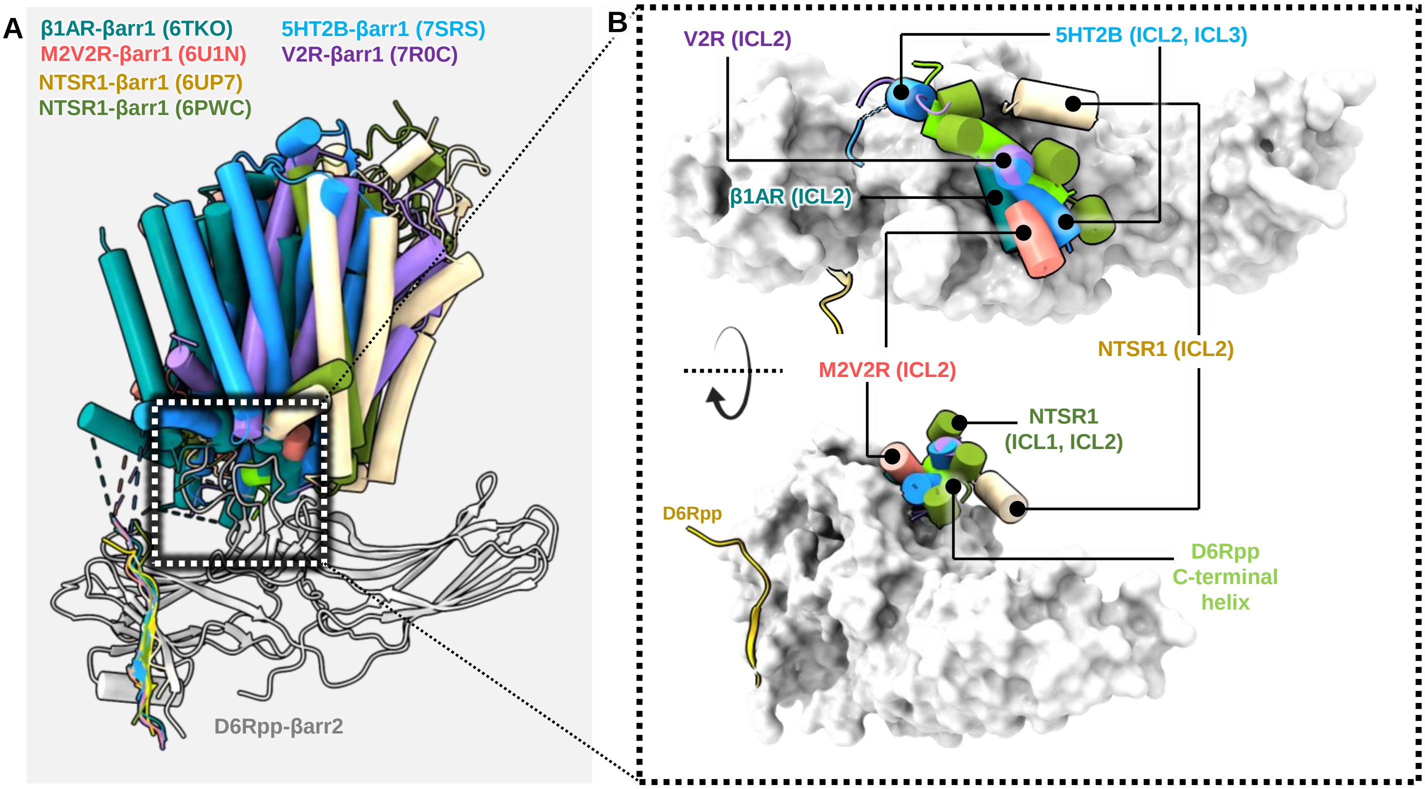

A
β1AR-βarr1 (6TKO)
M2V2R-βarr1 (6U1N)
NTSR1-βarr1 (6UP7)
NTSR1-βarr1 (6PWC)
5HT2B-βarr1 (7SRS)
V2R-βarr1 (7R0C)
D6Rpp-βarr2
B
V2R (ICL2)
5HT2B (ICL2, ICL3)
β1AR (ICL2)
M2V2R (ICL2)
NTSR1 (ICL2)
NTSR1
(ICL1, ICL2)
D6Rpp
D6Rpp
C-terminal
helix

**Figure S21. Intra-cellular loops of GPCRs dock onto the same site as the C-terminal helix in βarr2.**

**(A)** Superimposition of GPCR-βarr structures onto the D6Rpp activated βarr2 structure. **(B)** Snapshots of βarr2 in surface representation reveal that ICLs of GPCRs dock into the central crest of βarrs, which is the same site in the D6Rpp-βarr2 structure where the C-terminal helix binds.

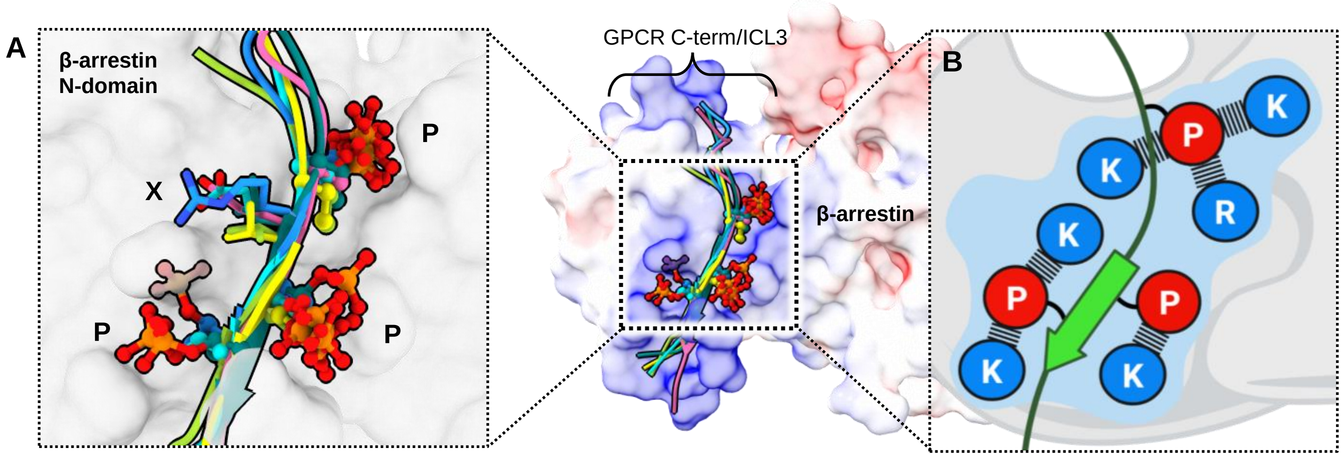
A
β-arrestin
N-domain
P
X
P
P
GPCR C-term/ICL3
β-arrestin
B
K
P
K
R
K
P
P
K
K

**Figure S22. Structural signature hallmarks of active βarr1/2.**

**(A)** The phosphorylated residues in GPCRs adopt a “PXPP” type motif for activation of βarrs. Structural alignment has been carried out on the highest resolution structure, M2Rpp-βarr2. (B) A cartoon representation of the conserved motifs, “PXPP” on 7TMRs and “KKRRKK” on βarrs have been illustrated.
